# Rapid, Low-Cost Detection of Water Contaminants Using Regulated *In Vitro* Transcription

**DOI:** 10.1101/619296

**Authors:** Khalid K. Alam, Jaeyoung K. Jung, Matthew S. Verosloff, Phillip R. Clauer, Jeong Wook Lee, Daiana A. Capdevila, Pablo A. Pastén, David P. Giedroc, James J. Collins, Julius B. Lucks

## Abstract

Synthetic biology has enabled the development of powerful nucleic acid diagnostic technologies for detecting pathogens and human health biomarkers. Here we expand the reach of synthetic biology-enabled diagnostics by developing a cell-free biosensing platform that uses <u>R</u>NA <u>o</u>utput <u>s</u>ensors <u>a</u>ctivated by ligand <u>ind</u>uction (ROSALIND) to detect harmful contaminants in aqueous samples. ROSALIND consists of three programmable components: highly-processive RNA polymerases, allosteric transcription factors, and synthetic DNA transcription templates. Together, these components allosterically regulate the *in vitro* transcription of a fluorescence-activating RNA aptamer: in the absence of a target compound, transcription is blocked, while in its presence a fluorescent signal is produced. We demonstrate that ROSALIND can be configured to detect a range of water contaminants, including antibiotics, toxic small molecules, and metals. Our cell-free biosensing platform, which can be freeze-dried for field deployment, creates a new capability for point-of-use monitoring of molecular species to address growing global crises in water quality and human health.

## INTRODUCTION

Synthetic biology is creating new capabilities for utilizing engineered biological systems to address pressing global challenges [1]. This is being achieved through uncovering fundamental engineering design principles for repurposing, evolving, and rewiring the functions of biological components for uses in a range of application contexts including biomanufacturing [2], bioremediation [3], and living medicines [4]. Among the most powerful and broadly useful functions of natural biological systems is the ability of cells to utilize molecular biosensors and genetic networks to sense and respond to changing internal and environmental conditions. This creates the powerful potential to harness natural biosensing capabilities for new diagnostic technologies that can vastly expand our ability to monitor human and environmental health [5].

This potential has started to be fulfilled with the development of synthetic biology-based nucleic acid diagnostics that can detect the presence of specific pathogens and human health biomarkers in complex matrices. These technologies utilize breakthroughs in RNA genetic switch design [6, 7] or CRISPR-mediated interactions [8, 9] to detect specific target DNA or RNA sequences. These molecular detection approaches are coupled to isothermal amplification strategies to enhance sensitivity, and genetically wired to the control of gene expression, fluorophore release, or other biochemical reactions to produce visual outputs when target molecules are present. The ability to freeze-dry reaction components in tubes or on paper for simple storage and distribution has led to breakthroughs in field-deployable diagnostic strategies for profiling the gut microbiome [10] and detecting human health pathogens such as Ebola, Zika and Norovirus [11–13] and plant pathogens such as Cucumber Mosaic Virus [14].

There is a tremendous need to extend the capability of synthetic biology diagnostics to sense chemical targets beyond nucleic acids. This is especially true in the case of water quality monitoring. Nearly 80% of the world’s population are at risk of water insecurity, partly due to a lack of access to clean and safely managed water [15]; yet clean water is essential for human activities including drinking, cooking, and agriculture. Among the many contaminants present in water, chemical species pose some of the most serious threats since they cannot be seen or easily detected. In fact, reliable methods to test water quality remain largely constrained to sophisticated, centralized laboratories with costly instrumentation that is time consuming and requires significant technical expertise. These limitations preclude implementation of water quality testing in the field, most notably in low-resource settings where they are most needed.

Here we address this need by developing a simple, low-cost synthetic biology platform using <u>R</u>NA <u>o</u>utput <u>s</u>ensors <u>a</u>ctivated by ligand <u>ind</u>uction (ROSALIND) to detect harmful contaminants in aqueous samples (**Fig. 1**). ROSALIND consists of three core components: (1) highly processive phage RNA polymerases, (2) allosteric transcription factors (aTFs), and (3) engineered DNA transcription templates, that together generate visible outputs upon exposure to specific ligands. The use of RNA-level outputs allows signals to be observed within minutes and addresses a key limitation of other cell-free approaches that rely on complex and resource-intensive protein translation to create detectable outputs. In addition, the use of simple and defined *in vitro* transcription (IVT) reactions overcomes a key limitation of more traditional whole-cell biosensing approaches by removing the need for growing and maintaining living cells, as well as the biocontainment and regulatory concerns of deploying genetically-modified organisms in the field.

**Fig. 1.**
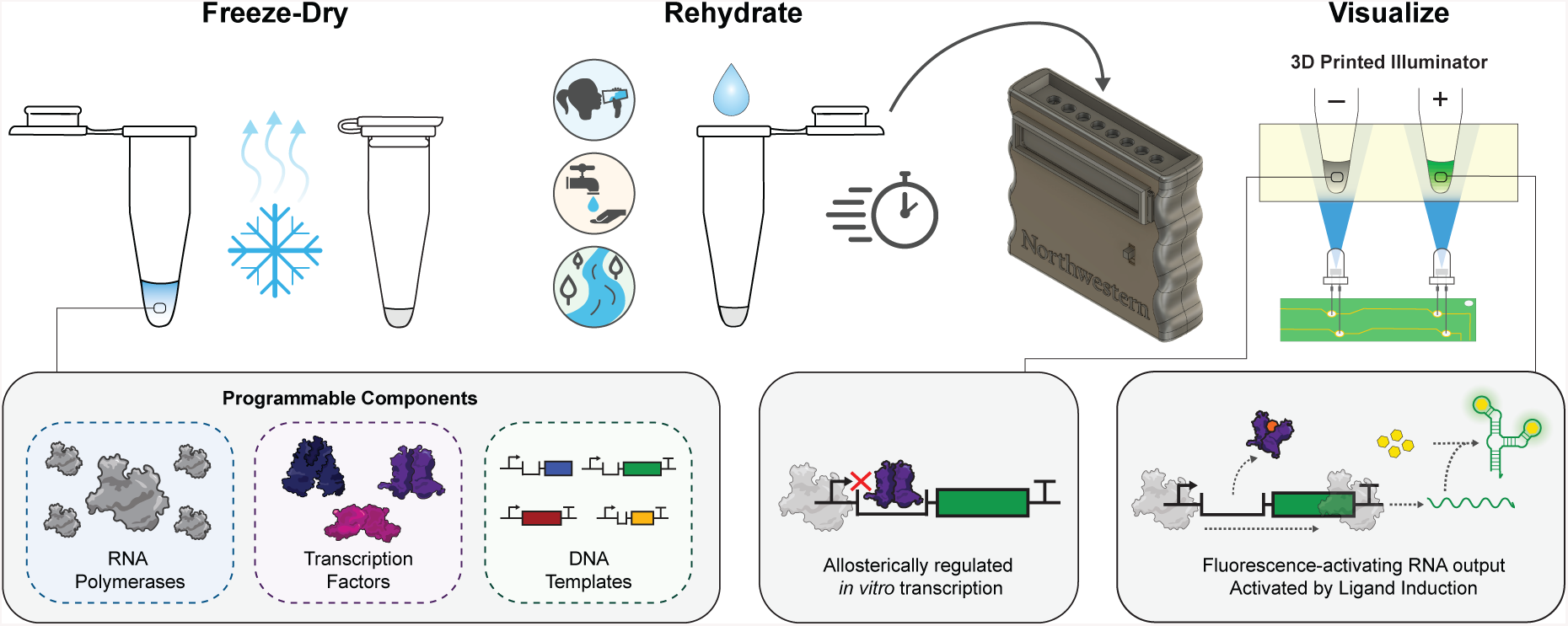
The ROSALIND platform. The RNA Output Sensor Activated by Ligand Induction (ROSALIND) platform consists of three programmable components: highly-processive RNA polymerases, allosteric transcription factors, and synthetic DNA transcription templates. Together, these components allosterically regulate the *in vitro* transcription of a fluorescence-activating RNA aptamer: in the absence of an analyte of interest, transcription is blocked, while in its presence a fluorescent signal is produced. ROSALIND can be freeze-dried for field deployment, thereby creating one drop water quality sensors that are activated upon rehydration with the sample of interest. A low-cost, 3D-printed handheld device provides easy visualization of the sensor’s RNA output.

We start by defining design rules for optimizing aTF-regulated IVT reactions using fast phage polymerases and show how the addition of specific chemical ligands can activate ROSALIND reactions to produce signals in tens of minutes. Our development of a simple 3D-printed handheld illuminator facilitates the visualization of ROSALIND outputs by eye. We next show that transcriptional outputs are compatible with RNA-level genetic circuitry to invert the response of repressive aTFs into activating systems more amenable to diagnostics. We showcase the modularity of ROSALIND through the demonstration of 16 different sensors that can detect compounds known or being investigated for their toxicity, including antibiotics, pharmaceuticals and personal care products, and heavy metals. We additionally report strategies to tune the threshold concentrations of target compounds above which ROSALIND produces visible output signals. Finally, we show that ROSALIND reactions can be freeze-dried for simple storage and distribution, and highlight an application of ROSALIND by testing environmental water samples known to be contaminated with copper above the United States Environmental Protection Agency (US EPA)-regulated drinking water threshold of 1.3 parts per million (ppm). As we highlight, ROSALIND represents a new capability for simple and rapid point-of-use monitoring of chemical species to address growing global health crises in water quality and human health.

## RESULTS

### *In vitro* transcription of a fluorescence-activating aptamer produces a fast and visible RNA output

A limitation of existing biosensing approaches is the use of protein-level outputs that must be transcribed and then translated through intricate and resource-intensive gene expression processes that lengthen the time to observation [16]. Additionally, protein-level reporters typically require either fluorophore maturation or significant enzymatic product accumulation prior to detection. We therefore reasoned that utilizing RNA-level outputs would allow biosensing reactions to be greatly simplified, leading to faster signal generation. To demonstrate this, we focused on fluorescence-activating RNA aptamers that bind and activate the fluorescence of otherwise non-fluorescent dyes [17, 18]. These aptamers have been engineered into allosteric biosensors for the *in vivo* detection of metabolites [19, 20] and as real-time reporters for monitoring IVT [21, 22], but have yet to be configured within transcriptional biosensing circuits as fast readouts of sensing events.

To demonstrate the viability of an RNA output sensor, we chose the three-way junction dimeric Broccoli (3WJdB) aptamer [23], an engineered variant that scaffolds two monomers of Broccoli around a highly structured RNA three-way junction motif, and the fast phage RNA polymerase (RNAP) from T7 bacteriophage. T7 RNAP, unlike *E. coli* or other native host RNA polymerases, is a highly processive, single-subunit enzyme used in biotechnological applications that require high levels of target gene expression. It is also extremely fast, with characterized transcription rates of 300 nucleotides (nt)/s *in vitro* [24]. To test the ability of T7 RNAP to drive 3WJdB synthesis in real-time, we PCR amplified a linear double-stranded DNA consisting of the T7 RNAP promoter followed by the 155 nt 3WJdB coding sequence and an optional 47 nt T7 transcription terminator [23] (**Fig. 2a**). T7 RNAP IVT was then performed using a commercially available transcription kit optimized for high yields of RNA and was supplemented with Broccoli-binding dye DFHBI-1T. Fluorescence activation was monitored in real-time and standardized to known amounts of fluorescein isothiocyanate (FITC), measured using the same excitation and emission wavelengths to make meaningful comparisons across experiments (**Supp. Fig. 1**). The IVT reactions generated micromolar amounts of mean equivalent FITC fluorescence (MEF) across a range of input DNA template concentrations within minutes (**Fig. 2b, Supp. Vid.**), illustrating the speed and viability of RNA-level outputs for biosensing applications. Furthermore, the fluorescence activation of 3WJdB was easily visualized at 0.5 MEF (µM FITC) with low-cost LEDs and a stage-lighting filter (**Fig. 2c**). Interestingly, we found that the inclusion of a T7 terminator at the end of the transcription templates resulted in improved fluorescence activation across the entire range of template concentrations used (**Supp. Fig. 2**).

**Fig. 2.**
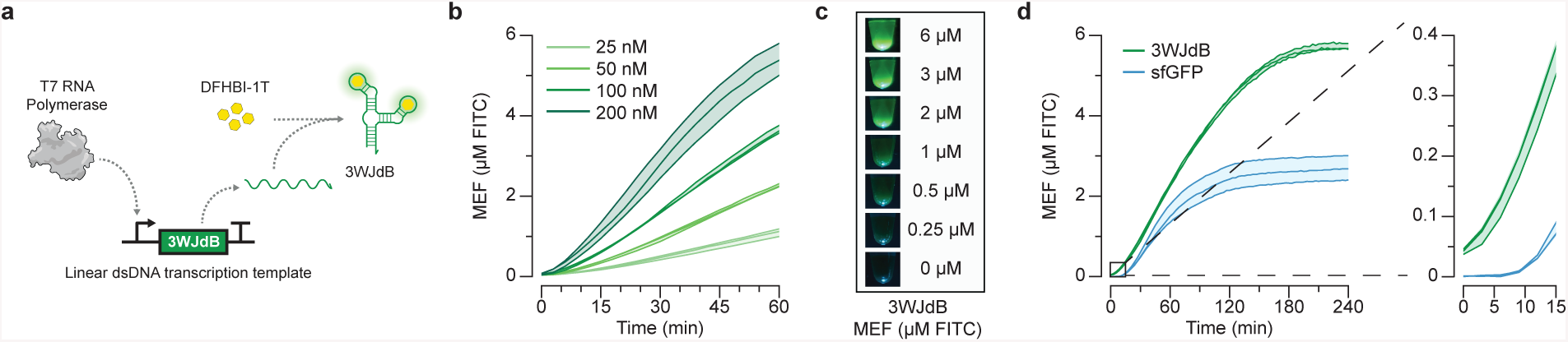
*In vitro* transcription of a fluorescence-activating aptamer rapidly generates a visible RNA output. **a**, T7 RNA polymerase transcription of the fluorescence-activating three-way junction dimeric Broccoli (3WJdB) aptamer. Binding of DFHBI-1T by 3WJdB activates its fluorescence. **b**, Titration of the linear dsDNA transcription template in a commercially-available transcription kit generates micromolar amounts of mean equivalent fluorescence (MEF) when standardized to soluble fluorescein isothiocyanate (µM FITC) measured under identical plate reader conditions. **c**, Visual detection of 3WJdB:DFHBI-1T fluorescence, with values corresponding to MEF of soluble FITC. **d**, Comparison of fluorescence kinetics using an equimolar concentration of DNA template (50 nM) encoding either 3WJdB in a commercially available transcription kit or sfGFP in commercially-available transcription-translation kit. All data shown for 3 experimental replicates as individual lines with raw fluorescence values standardized to MEF (µM FITC). Shading indicates the average value ± standard deviation.

We next compared the fluorescence intensity and kinetics of the RNA output generated using a commercial transcription kit to that of superfolder GFP (sfGFP) [25], a protein-level reporter commonly utilized for its fast fluorophore maturation rate, produced using a commercially available transcription-and-translation kit based on the protein synthesis using recombinant elements (PURE) system [26]. Protein expression in the PURE system is driven by T7 RNAP and coupled to translation of the resulting mRNA by reconstituted translational machinery, including ribosomes, elongation factors, and tRNA synthetases. It is widely used in emerging cell-free biotechnologies [27], including paper-based diagnostics [12], because of its defined reaction composition. 3WJdB-expressing IVT reactions and sfGFP-expressing PURE reactions were set up and activated using equimolar amounts of DNA templates. In this setup, the RNA output immediately began to increase in fluorescence intensity, while sfGFP required at least nine minutes before a detectable increase in signal was observed (**Fig. 2d**). This remained true across a range of input template concentrations (**Supp. Fig. 3**). Additionally, over the entire four-hour time course, 3WJdB transcription produced substantially more fluorescence than sfGFP. Finally, we compared the commercial transcription kit to a homemade transcription reaction using in-house purified T7 RNAP. We found that the in-house reactions performed as well as the commercial kit over a one-hour time course (**Supp. Fig. 4**), had an advantage of being of known composition compared to the commercial kit, and offered a significant reduction of the cost per reaction to <1 USD at laboratory scale (**Supp. Table 1**), representing a fifth of the cost of a commercial IVT kit and a twenty-fifth of the cost of the PURE kit.

Together, these results demonstrate that RNA-level outputs generated from simple IVT reactions represent a viable and even advantageous approach for biosensing. By eliminating the need for protein synthesis, our transcriptional sensing approach significantly reduces the cost and complexity associated with generating a visibly fluorescent output.

### *In vitro* transcriptions can be configured into ligand-responsive biosensors using allosteric transcription factors

We next sought to create a ligand-responsive biosensor by using an aTF to regulate T7 RNAP IVT. We reasoned that the binding of an aTF to its cognate operator sequence would prevent T7 RNAP from switching from initiation to elongation phase in the absence of ligand, and ligand-mediated derepression of an aTF would result in a fluorescence signal (**Fig. 1**).

To implement this design, we first investigated how aTF-operator interactions can block T7 RNAP transcription in the absence of ligand. We designed synthetic transcription templates that introduced the aTF operator sequence at an appropriate distance from the promoter sequence, such that aTF binding would prevent the conformational transition of T7 RNAP from the highly-abortive initiation phase of transcription to the productive elongation phase, which typically occurs after 8-12 nts have been transcribed [28, 29]. We chose TetR as a model aTF to test these designs. TetR is a well-characterized aTF that regulates the tetracycline efflux pump encoded by *tetA* and assumes a two-domain architecture comprising a three-helix bundle DNA-binding domain and a ligand-binding/dimerization domain [30]. It binds to a 19 base-pair (BP) palindromic *tet* operator sequence (*tetO*) as a 46 kDa dimer with low nanomolar affinity [31]. Furthermore, the TetR-*tetO* interaction is disrupted in the presence of tetracyclines, which bind to TetR at the dimerization interface and trigger an allosteric change that substantially reduces TetR’s affinity for *tetO* [31]. We designed a series of transcription templates to explore the effect of spacing between the minimal 17 BP T7 promoter and the downstream *tetO* sequence (**Fig. 3a**). We utilized the native sequence that follows the canonical T7 RNAP promoter (Class III Φ10) [32–34] as a spacer in two BP increments from 0-14 BP. Immediately following the spacer, we placed the *tetO* sequence, followed by 3WJdB and the T7 terminator. IVT reactions were setup as before for the in-house reactions, with the addition of purified recombinant TetR protein in 100-fold excess of the DNA transcription template (2.5 µM dimeric TetR, 25 nM DNA). We observed that the absence of any spacing (0 BP) resulted in no fluorescence activation in either the presence (regulated) or absence (unregulated) of TetR, likely due to the lack of the strongly preferred G base at the T7 RNAP transcription start site [35] (**Fig. 3b**). However, robust fluorescence activation was observed without TetR when using a two BP spacer, which was reduced by 100-fold to baseline levels when regulated by TetR. The trend of strong fluorescence activation in the unregulated reaction and strong repression in the regulated reaction was consistent over a two to six BP spacer length, whereas fluorescence activation in the regulated reactions increased for spacers that were eight BP or longer. In particular, over a four-hour time course, the use of a four BP spacer in the presence of TetR resulted in >100-fold repression without any background subtraction, representing virtually no transcription leak (**Fig. 3c**). We therefore chose these shorter spacers for subsequent promoter-operator designs, as this ensured both productive transcription by using the preferred initiating nucleotides and strong repression in the presence of the aTF.

**Fig. 3.**
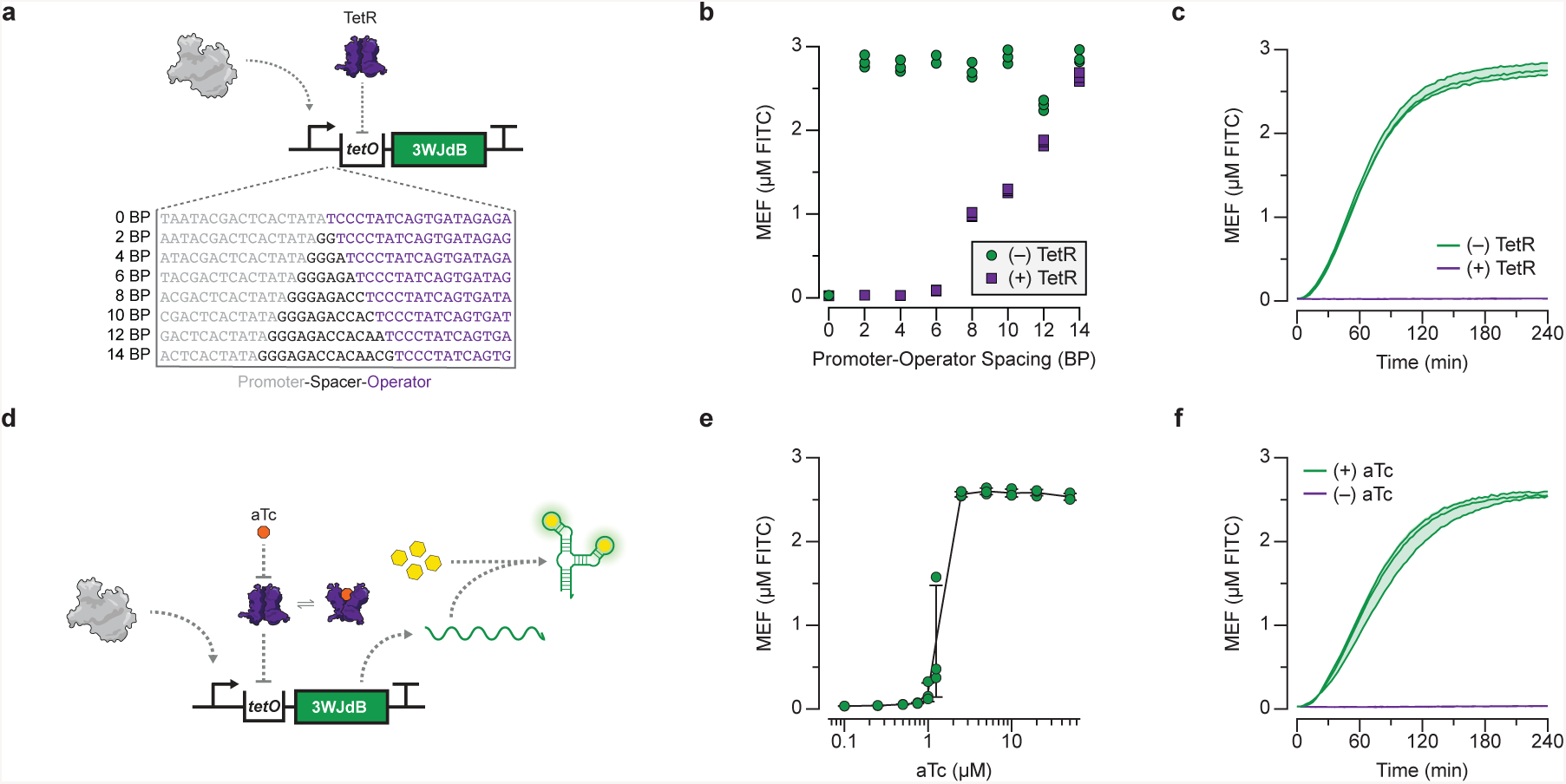
Allosteric regulation of *in vitro* transcription. *In vitro* transcriptions can be allosterically regulated by designing a synthetic transcription template configured to bind a purified transcription factor (TetR) via the operator sequence (*tetO*) placed downstream of the T7 promoter. When bound by TetR, the aTF serves as a steric block to transcription elongation. **a**, Spacing between the T7 promoter and *tet*O site influences the ability to regulate transcription in the presence of TetR. A series of spacers in 2 base pair (BP) intervals was constructed to evaluate the impact of spacer length on the ability to regulate transcription. **b**, End point data shown for promoter-operator spacer variants regulated (with 2.5 µM TetR dimer, 100-fold excess over DNA template) and unregulated (without TetR) after 240 minutes. **c**, Kinetic comparison of 3WJdB fluorescence activation of TetR-regulated versus unregulated transcription reactions with a 4 BP spacer. **d**, Induction of a TetR regulated *in vitro* transcription reaction occurs in the presence of the cognate ligand, anhydrotetracycline (aTc), by binding to TetR and preventing its binding to the DNA template. This allows transcription to proceed, leading to fluorescence activation. **e**, Dose response with anhydrotetracycline (aTc), measured at 240 minutes with 25 nM DNA and 1.25 µM TetR dimer. **f**, Kinetics of induction (from **e**) with 2.5 µM aTc. All data shown for 3 experimental replicates as lines or points with raw fluorescence values standardized to MEF (µM FITC). Shading (**c, f**) and error bars (**e**) indicate the average value of experimental replicates ± standard deviation.

We next sought to determine if TetR can be derepressed with a cognate ligand to permit IVT and therefore act as a biosensor (**Fig. 3d**). We first performed IVT reactions over a range of TetR concentrations. We found that a 25-fold excess of TetR dimer (625 nM) to transcription template (25 nM) was the minimal amount of repressor needed to ensure tight regulation of transcription, while a 50-fold excess of TetR dimer (1.25 µM) performed similarly to the 100-fold excess that effectively repressed all fluorescence activation (**Supp. Fig. 5**). We then setup IVT reactions using these TetR and DNA template concentrations (25 nM DNA, 1.25 µM TetR dimer), and included a range of anhydrotetracycline (aTc) concentrations, a high-affinity ligand of TetR often used in synthetic biology applications due to its non-antibiotic properties when compared to other tetracyclines. We observe strong repression over four hours through high nanomolar amounts of aTc, with half-maximal induction between 1 and 2.5 µM aTc (**Fig. 3e**). Importantly, when induced with 2.5 µM aTc, the kinetics of induction follow those of the unregulated transcription reaction (**Fig. 3c,f**).

**Fig. 4.**
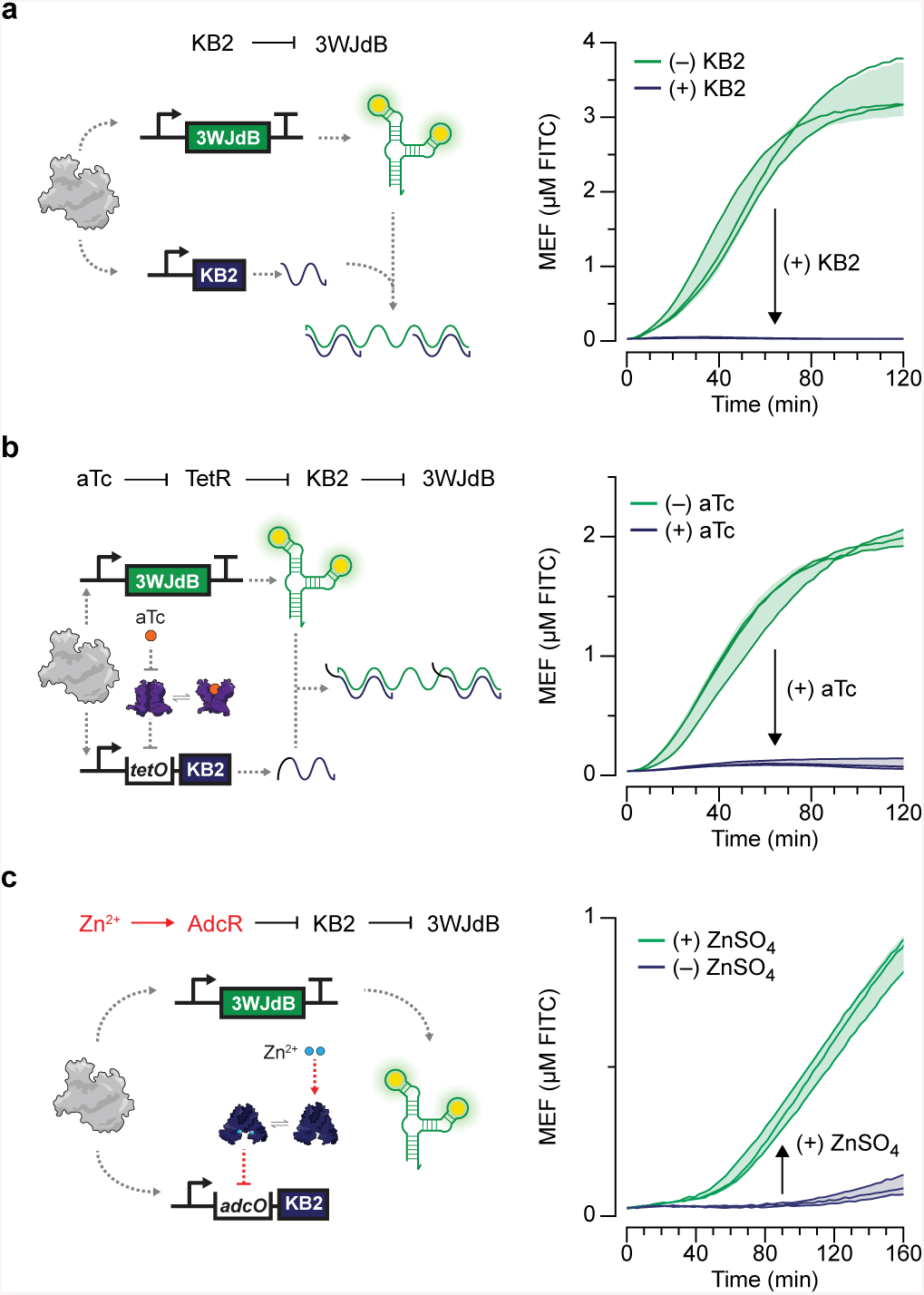
Transcription factor responses can be inverted with RNA kleptamers to expand the scope of ROSALIND-compatible transcription factors. **a**, A kleptamer RNA strand (KB2) antisense to the dye-binding region of the broccoli aptamer can disrupt folding and lead to the loss of fluorescence. Addition of a KB2-expressing template in a 4:1 ratio with the 3WJdB template inhibits signal. **b**, Kleptamers can be used to invert the response of transcription factors. KB2 transcription can be regulated by TetR (1.25 μM dimer) and aTc (2.5 μM shown) by placing the *tetO* site in between the T7 promoter and KB2 coding sequence. **c**, This scheme was used to create a ROSALIND zinc sensor using the aporepressor AdcR. When bound to Zn^2+^ (30 μM), AdcR (1.5 μM dimer) binds to its cognate operator sequence, *adcO*, placed upstream of the KB2 coding sequence, preventing KB2 from being expressed and thereby activating fluorescence. Arrows inside of the plots represent direction of regulation when indicated species are added. All data shown for 3 experimental replicates as lines with raw fluorescence values standardized to MEF (µM FITC). Shading indicates the average value ± standard deviation. 3WJdB DNA template concentrations used are: 25 nM for **a, b** and 7.5 nM for **c**. KB2 DNA template concentrations used are: 100 nM for **a**, and 150 nM for **b, c**.

**Fig. 5.**
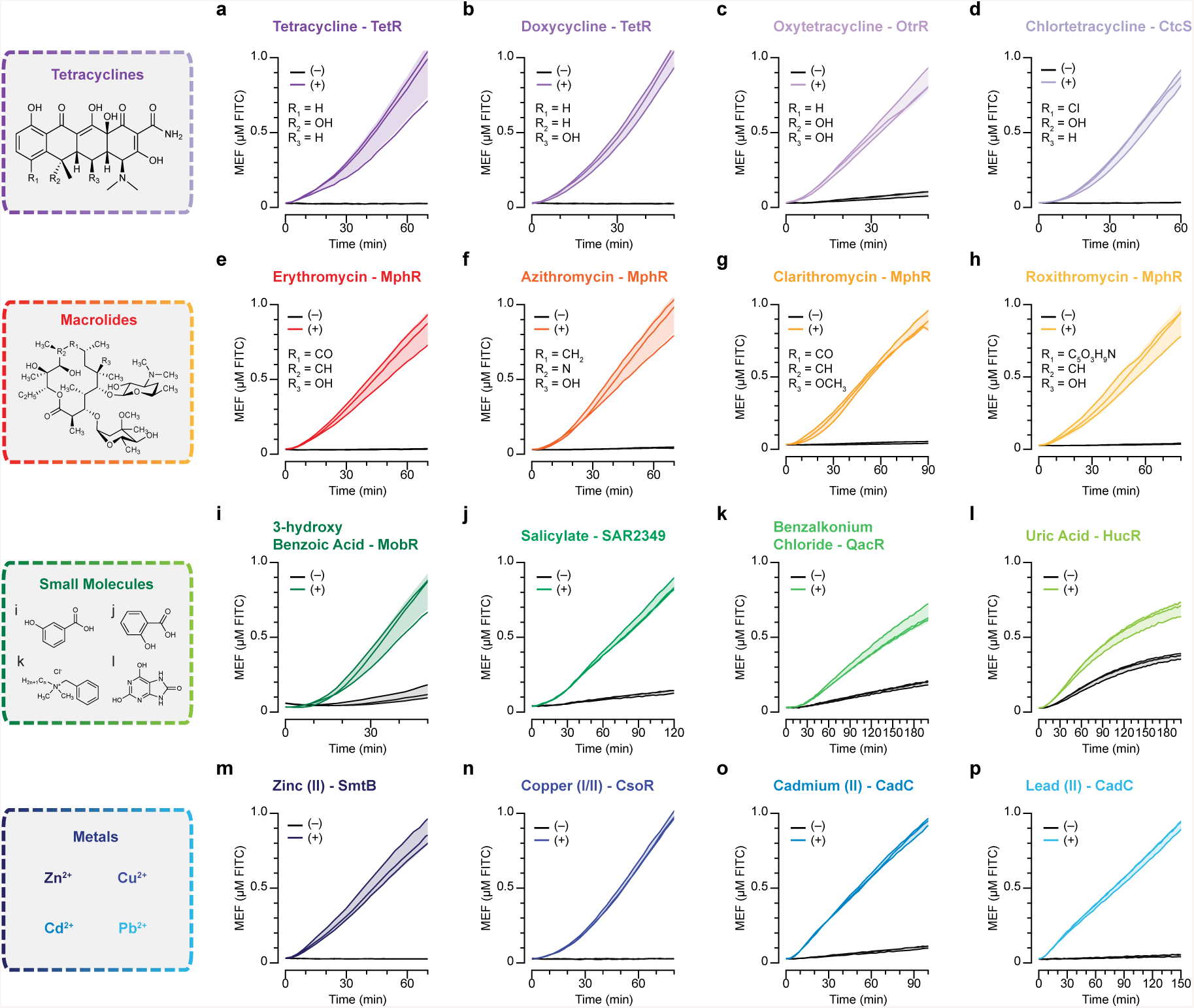
ROSALIND is modular and can be used to sense complex molecules, small molecules and metal ions. TetR (1.25 μM dimer) can be used to sense **a**, tetracycline (25 μM), **b**, doxycycline (12.5 μM) and other tetracycline family antibiotics. **c**, Oxytetracycline sensing (100 μM) using OtrR (2.5 μM dimer), a MarR-family aTF from the oxytetracycline biosynthesis gene cluster from *Streptomyces rimosus*. **d**, Chlortetracycline sensing (50 μM) with CtcS (1.25 μM dimer), a MarR-family aTF from the chlortetracycline biosynthesis gene cluster from *Streptomyces aureofaciens*. MphR (0.625 μM dimer), a TetR-family aTF from the *Escherichia coli* macrolide resistance mechanism senses the macrolides **e**, erythromycin (50 μM), **f**, azithromycin (50 μM), **g**, clarithromycin (50 μM) and **h**, roxithromycin (250 μM)**. i**, 3-hydroxybenzoate sensing (2 mM) with MobR (100 μM dimer), a MarR-family aTF from *Comamonas testosteroni* KH122-3S. **j**, Salicylate sensing (12 mM) using SAR2349 (50 μM dimer), a MarR-family aTF from *Staphylococcus aureus*. **k**, Benzalkonium chloride (100 μM) sensing with QacR (5 μM dimer), a TetR-family aTF from *Staphylococcus aureus* that responds to quaternary ammonium cations. **l**, Uric acid (303 μM) sensing using HucR (2.15 μM dimer), a MarR-family aTF from *Deinococcus radiodurans*. **m**, Zn^2+^ sensing (10 μM ZnCl_2_) with SmtB (5 μM dimer), a ArsR/SmtB-family aTF from *Synechococcus elongatus* PCC 7942. **n**, Cu^2+^/Cu^+^ sensing (10 μM CuSO_4_ in 10 mM DTT) with CsoR (2.5 μM tetramer), an aTF from *Bacillus subtilis***. o**, Cd^2+^ sensing (10 μM CdCl_2_) and **p**, Pb^2+^ sensing (51.8 μM PbCl_2_) with the ArsR/SmtB family aTF CadC (1.5 μM dimer), a SmtB/ArsR family aTF from *Staphylococcus aureus* pl258. Each reaction contains the indicated amount of ligand (+) dissolved in laboratory-grade water, ethanol, Tris-base buffer, or diluted sodium hydroxide, or a laboratory-grade water control (-) (**Supp. Table 3**). All data shown for 3 experimental replicates as lines with raw fluorescence values standardized to MEF (µM FITC). Shading indicates the average value ± standard deviation. DNA template concentrations used in each reaction are 25 nM for **a-h, k, m, n**, 10 nM for **i, j, p**, 17.5 nM for **l**, and 20 nM for **o**.

Together, these results show that aTF-regulated IVT of fluorescence-activating RNA aptamers can serve as fast and cheap biosensors, establishing a proof-of-concept basis for the ROSALIND platform.

### RNA circuitry can invert transcription factor responses *in vitro*

The above results show that aTFs can be used to regulate IVT in response to ligands through derepression of operator binding. However, there are aTFs, often referred to as aporepressors, that function in the inverse manner—by binding the operator site only when bound to their cognate ligands [36–38]. While this ligand corepression represents an important control of gene expression, it is not as useful in a diagnostic biosensor context where direct activation of visible outputs upon exposure to target ligands is often desired. To address this, we sought to develop a strategy to invert the response from aporepressors by manipulating the transcriptional outputs with RNA genetic circuitry.

To achieve this, we leveraged the recent development of RNA kleptamers—short antisense RNAs designed to be complementary to the dye-binding region of the Broccoli aptamer [39]. When present, the KB2 kleptamer inhibits Broccoli fluorescence by binding to and disrupting the aptamer from folding into the conformation required for fluorescence activation. We thus reasoned that we could invert the output of an aTF-regulated transcription by using the aTF to regulate KB2 expression, which would then interfere with a constitutively expressed 3WJdB.

We first examined if the fluorescence activation of the 3WJdB aptamer could be inhibited through unregulated IVT of KB2 with T7 RNAP. Titration of the KB2 DNA template in IVT reactions resulted in the proportional decrease of fluorescence activation (**Supp. Fig. 6a**), and a 4:1 ratio of the KB2 to 3WJdB transcription templates resulted in complete inhibition of fluorescence (**Fig. 4a**). We then sought to regulate the transcription of KB2 with TetR, by placing the *tetO* sequence five BP downstream of the T7 promoter and immediately upstream of the KB2 sequence. Similar to the direct repression of 3WJdB transcription (**Fig. 3c**), TetR was able to repress KB2 transcription at 2.5 µM or above, resulting in fluorescence activation from the constitutively expressed 3WJdB (**Supp. Fig. 6b**). Addition of its cognate ligand, aTc, at 2.5 µM or above effectively derepressed TetR from the KB2 template, thereby allowing KB2 transcription and resulting in switching off the fluorescence of 3WJdB (**Fig. 4b**).

**Fig. 6.**
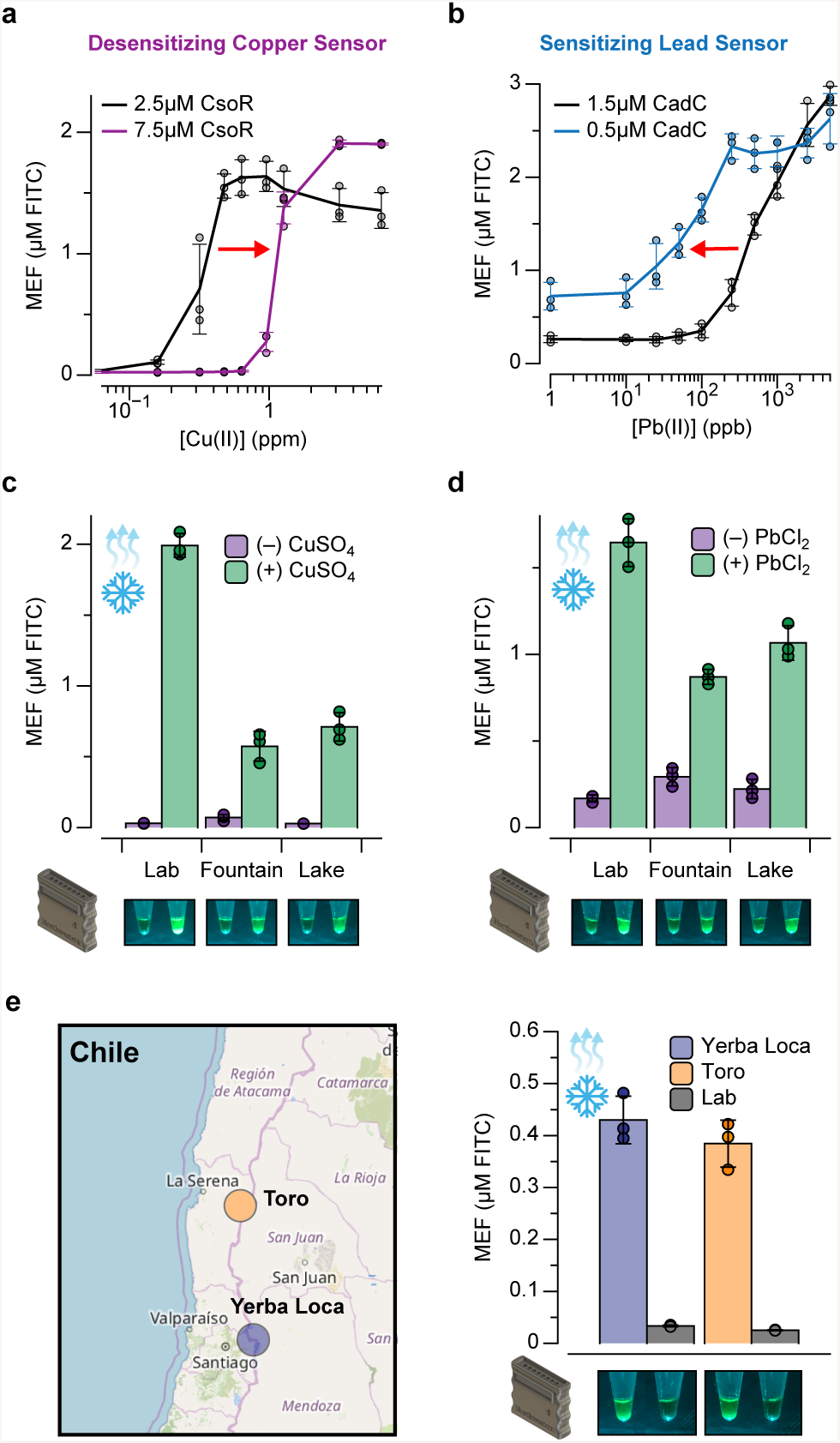
ROSALIND can be optimized for field deployment. ROSALIND sensitivity can be tuned by changing the concentration of transcription factor. **a**, Increasing the concentration of CsoR desensitizes the copper sensor (dose response shift to higher concentrations), while **b**, decreasing the concentration of CadC sensitizes the lead sensor (dose response shift to lower concentrations). The red arrows in the plots indicate the direction of dose response shift. ROSALIND reactions can be freeze-dried and rehydrated with various water sources. Reactions are lyophilized overnight and rehydrated with laboratory-grade water, drinking water (fountain), and lake water spiked with **c**, 0.0 ppm and 1.3 ppm of CuSO_4_ and **d**, 0.0 ppm and 2 ppm of PbCl_2_. Fluorescence results shown after 4 hours of incubation at 37°C with plate reader quantification (top) and hand-held illuminator images (bottom). ROSALIND can detect contaminated levels of metals in environmental water samples. **e**, Water samples from the Yerba Loca Creek and the Toro River, Chile, contaminated with >7 ppm of copper were tested with freeze-dried copper sensors. Water samples were filtered to remove particulate matter, and then serial diluted (1:8 dilution shown). The diluted samples were independently validated to contain 0.88 ppm (Yerba Loca) and 1.05 ppm (Toro) copper with flame atomic absorption spectroscopy. All data shown for 3 experimental replicates as points with raw fluorescence values standardized to MEF (µM FITC). Bars represent averages over three replicates and error bars represent ± standard deviations. DNA template concentrations used are 25 nM except for the 0.5 μM CadC data which has 10 nM DNA template. The map shown in **e** was provided by courtesy of OpenStreetMap contributors.

To test whether this strategy could be implemented to use aporepressors in a diagnostic biosensor context, we applied it to AdcR, a metal-sensing member of the MarR-type transcription factor family from *Streptococcus pneumoniae* [40, 41]. In its native context, AdcR in the presence of Zn(II) downregulates the expression of *adcCBA*, a high-affinity ABC uptake system for Zn(II), and genes encoding cell-surface Zn(II)-binding proteins. This is mediated by two Zn(II) atoms which bind to a homodimer of AdcR as corepressors and enable the complex to bind and repress its operator sequence (*adcO*). Similar to the TetR-regulated KB2, we placed *adcO* five BP downstream of the T7 promoter sequence and immediately upstream of the KB2 sequence. The addition of 1.5 µM AdcR dimer alone was unable to repress the KB2 template, and therefore no fluorescence was observed. However, fluorescence activation was observed when 30 µM of Zn(II) was supplemented into the reaction (**Fig. 4c**). Overall, these results demonstrate that aporepressors can be reconfigured to exhibit ROSALIND-compatible ligand-activated behavior using RNA genetic circuitry.

### ROSALIND can be modularly reconfigured to sense antibiotics, small molecules and metal ions

Having shown that aTFs can be rewired to control transcription in two different contexts, we next sought to expand the range of molecules that can be sensed with the ROSALIND platform by leveraging the natural diversity of aTFs. Natural aTFs have evolved to sense a variety of complex molecules, small molecules, and heavy metals [38]. We therefore reasoned that ROSALIND could be configured to sense this range of important target compounds and elements simply by modularly changing the aTF-operator pair within the transcription templates and including the corresponding aTF in the reaction.

We began by focusing on antibiotic detection. Overuse of antibiotics in human and veterinary medicine, agriculture, and aquaculture is resulting in significant contamination of freshwater supplies [42]. The presence of antibiotics at sublethal concentrations in these settings drives antimicrobial resistance [43], and rapid and frequent monitoring of antibiotics is important to prevent the emergence of multi-drug resistant bacteria. We first tested our TetR-based sensor to detect several members of the tetracycline family of antibiotics, including tetracycline (**Fig. 5a**) and the semi-synthetic antibiotic doxycycline (**Fig. 5b**), both of which are on the World Health Organization’s list of essential medicines. To test the ability of ROSALIND to sense these antibiotics, TetR-regulated transcription templates were included in reactions that contained purified TetR; reactions were run using either laboratory-grade water (-), or similar water with added antibiotics (+). In both cases, we observed clear generation of fluorescence with the antibiotic present and no fluorescence in the purified water control.

Following this procedure, we next expanded beyond TetR-sensing of tetracyclines with OtrR, a recently characterized aTF that regulates the oxytetracycline biosynthetic gene cluster from *Streptomyces rimosus* [44]. OtrR is a member of the multiple antibiotic resistance repressor (MarR) family of aTFs [40], and has previously been used to create a whole-cell biosensor for oxytetracycline [45]. When applied within ROSALIND, OtrR and its corresponding DNA template were capable of acting as a biosensor for oxytetracycline (**Fig. 5c**).

To expand the range of tetracycline antibiotic sensing even further, we identified a putative MarR aTF, CtcS, found in the chlortetracycline biosynthetic gene cluster of an industrial strain of *Streptomyces aureofaciens* [46]. Using knowledge of the natural context of MarR aTFs, we identified the divergent operon that controls both the expression of the chlortetracycline efflux pump and the putative aTF analogous to OtrR and designed a minimal operator sequence for CtcS to enable its use in ROSALIND. As with OtrR, CtcS and its corresponding DNA template were capable of acting as a biosensor for chlortetracycline (**Fig. 5d**). As this is the first *in vitro* characterization of CtcS, this result demonstrates the potential of ROSALIND as a screening and characterization platform for aTFs.

We next sought to expand sensing to another class of antibiotics that are also widely used in human health, namely, macrolides. We therefore chose MphR, a TetR family aTF used in *E. coli* to control expression of *mphA*, a gene encoding a macrolide 2’- phosphotransferase that confers resistance to macrolide antibiotics through inactivation [47]. MphR has previously been shown to respond to macrolides such as erythromycin, a drinking water contaminant of emerging concern [48], and has been engineered into whole-cell macrolide biosensors [49, 50]. When configured in ROSALIND, MphR and its corresponding DNA template were able to detect a family of macrolide antibiotics, including erythromycin, azithromycin, clarithromycin and roxithromycin (**Fig. 5e-h**). We note that this observed broad range of MphR specificity is expected based on its known promiscuity and is likely a common feature of antibiotic-sensing aTFs [49, 50].

Having demonstrated the ability of ROSALIND to detect complex antibiotic molecules, we next sought to leverage the platform for the detection of small molecules, specifically focusing on ones that can be environmental contaminants or human heath markers. In particular, there has been a growing concern that non-regulated contaminants, such as those generated from the large use of pharmaceuticals and personal care products (PPCPs), interfere with or are unaffected by water treatment, and are causing widespread, but poorly characterized health and environmental concerns [51]. Since these contaminants of emerging concern can only be detected using sophisticated laboratory equipment, there is a considerable and growing need for rapid, low-cost sensing platforms that can be used to assess the presence of PPCP contamination in water supplies. As a model PPCP, we started with 3-hydroxy benzoic acid, a perfume additive that can induce the expression of hydroxybenzoate hydroxylase by derepressing the MarR-family aTF MobR within the aerobic soil bacterium *Comamonas testosteroni* KH122-3S [52]. When ROSALIND reactions were configured with MobR and its corresponding DNA template, we again observed strong activation of fluorescence only in the presence of 3-hydroxy benzoic acid (**Fig. 5i**), indicating that the ROSALIND platform can be configured to sense small molecules of interest.

We next sought to demonstrate that ROSALIND could be used to sense two PPCP compounds: (1) salicylate, a widely used compound in personal care products [48] that has been shown to bind to the *Staphylococcus aureus* MarR-family aTF SAR2349 [53], and (2) benzalkonium chloride, a widely used disinfectant that can promote antimicrobial resistance [54], and has been shown to bind to the *Staphylococcus aureus* TetR-family aTF QacR [55]. When these purified aTFs and the corresponding operator-containing DNA constructs were included in ROSALIND reactions, we were able detect these two PPCP compounds at a high micromolar to low millimolar range (**Fig. 5j**,**k**).

In addition to being chemical contaminants in water, small molecules can also be important human health biomarkers. For example, the accumulation of uric acid can be used as an early indicator of gout when present at 6 mg/dL or above within bodily fluids [56]. As a proof of concept, we thus configured ROSALIND reactions to sense uric acid by incorporating purified HucR, a MarR-family aTF from *Deinococcus radiodurans* [57], and the corresponding DNA template, and showed that we could detect uric acid within water samples approximately at 300 µM (∼5 mg/dL) (**Fig. 5l**).

Among water contaminants, metal contamination in drinking water has recently come to the forefront as a major issue. Specifically, alarming amounts of copper, lead, cadmium, arsenic, mercury and other metals are being discovered in drinking water sources across the globe due to aging infrastructure, environmental disasters, industrial and consumer waste, and from natural abundance in groundwater [58–60]. The current state-of-the-art for metal detection in water requires traditional chemical analysis in centralized laboratories, which can be slow, costly, and require significant technical expertise—all of which prohibit consumers from easily monitoring their water supplies. We thus reasoned that, properly configured, ROSALIND could provide a new capability for the rapid detection of metals in water that would address a key need for global public health. As a proof-of-concept we identified an additional zinc sensing aTF, SmtB from the ArsR/SmtB family of regulators [61, 62]. Unlike AdcR, SmtB acts as a repressor when Zn(II) is absent and therefore did not require the kleptamer inversion strategy for compatibility with ROSALIND. Using SmtB and the corresponding DNA template, we were able to sense Zn(II) at 10 µM (0.65 ppm), which is more sensitive than the US EPA secondary drinking guideline of 5 ppm (**Fig. 5m**).

While many metals are toxic at high concentrations, of particular concern are copper and lead [59, 60, 63]. Psychiatric disorders, impaired neural development, and organ damage can all result from chronic exposure to high levels of copper and lead. Of note, copper and lead levels are strictly regulated in drinking water by the US EPA’s Lead and Copper Rule (LCR). Among the many copper-sensing aTFs is CsoR, a Cu(I) sensing transcriptional regulator from *Mycobacterium tuberculosis* and other bacteria that forms a homotetramer to bind four copper ions [64, 65]. When configured into ROSALIND with the appropriate DNA template, our sensor was able to detect copper at 10 µM (0.6 ppm) (**Fig. 5n**). Interestingly, while the cognate ligand for CsoR is Cu(I), we observed that the reaction senses Cu(II) from added CuSO_4_. This could be the result of the presence of 10 mM DTT in IVT reactions that can serve to reduce Cu(II) to Cu(I). Another ArsR/SmtB family aTF, CadC, is found in pI258 of *Staphylococcus aureus* and known to bind a number of metals, including cadmium and lead [66]. When implemented in ROSALIND with appropriately configured DNA templates, we were able to robustly detect both cadmium and lead at 10 µM (1.12 ppm) and 51.8 µM (10.7 ppm), respectively (**Fig. 5o**,**p**).

In total, we employed 12 different aTFs (**Supp. Fig. 7**) to sense 17 different ligands that are structurally and functionally diverse. In most cases, a single aTF is reported to sense multiple ligands, as is the case with TetR sensing tetracycline derivatives, MphR sensing macrolide derivatives, and CadC sensing both lead and cadmium. In these cases, ROSALIND reactions configured with these aTFs and corresponding DNA templates are best viewed as reporting on the presence of a specified family of related compounds or elements. To investigate whether ROSALIND reactions are specific for the indicated compounds or elements outside of these examples, we chose tetracycline, erythromycin, salicylate and copper as representative members of each class of compounds/elements and challenged them against ROSALIND reactions containing TetR, MphR, SAR2349 and CsoR. We found strong signals from cognate ligands with minimal cross-talk indicating that ROSALIND reactions are specific for their intended ligands (**Supp. Fig. 8**). Together, these results show that the ROSALIND platform is highly modular and that aTFs function with their expected specificity.

### ROSALIND can be tuned and applied to real-world water samples

Application of ROSALIND to real-world water/environmental samples requires that sensitivity thresholds can be tuned to be near the relevant levels for each contaminant, and that the reactions can be formulated for distribution and use in the field. To explore whether ROSALIND can be tuned to meet these criteria, we chose to focus on copper sensing and lead sensing reactions using CsoR and CadC, respectively, as proof-of-concept demonstrations.

We first sought to develop strategies that can be used to tune the sensitivity of ROSALIND reactions. A titration curve of the CsoR-containing ROSALIND reaction showed that the sensor produces signals greater than 0.5 MEF (µM FITC), which we found to be visible by eye (**Fig. 2c**), at added CuSO_4_ concentrations of 5 µM (0.3 ppm of Cu(II)) and above. This indicates that the sensor was sensitive to levels far below the threshold set by the LCR (1.3 ppm) (**Fig. 6a**). To desensitize the sensor to respond near the LCR threshold, we reasoned that excess CsoR could be added to the reactions to act as a copper chelator. As expected, when CsoR tetramer concentration was increased from 2.5 µM to 7.5 µM—while keeping the DNA template concentration constant at 25 nM—the ROSALIND dose response curve was shifted and enabled reactions to activate near the desired threshold of 1.3 ppm of copper from added CuSO_4_ (**Fig. 6a**). For lead sensing, the challenge was the opposite—ROSALIND reactions configured with 1.5 µM of CadC dimer and 25 nM of the corresponding DNA template were activated at Pb(II) concentrations near 250 ppb, which was an order of magnitude higher than the accepted safety guidelines of 15 ppb (**Fig. 6b**). To sensitize the lead sensor, we thus employed the converse of our CsoR strategy. By lowering CadC dimer concentration to 0.5 µM, we were able to shift the dose response curve to be more sensitive in order to detect tens of ppb Pb(II) (**Fig. 6b**). These results demonstrate that ROSALIND sensitivity can be tuned by simply adjusting the amount of aTF and corresponding DNA template in the reactions.

We next examined whether ROSALIND can be formulated for distribution in the field. Many of the advances in synthetic biology diagnostic technologies have been enabled through the demonstration that cell-free systems can be freeze-dried for convenient storage and distribution before rehydration at the point-of-use [2, 11, 12]. However, it was unclear if similar strategies would work for ROSALIND, since it is a purely *in vitro* transcription system. To test this, we explored a range of known lyoprotectants and determined that nonreducing disaccharides, such as sucrose and trehalose, in combination with a sugar alcohol, mannitol, provided substantial lyoprotection. We applied these additives to the copper and lead sensors configured with 7.5 µM tetrameric CsoR (25 nM DNA) and 1.5 µM dimeric CadC (20 nM DNA), respectively. Freeze-dried copper and lead sensors were then tested by rehydrating with three different water sources: laboratory-grade water, drinking water sampled from a water fountain, and Lake Michigan water. Each water source was spiked with either 2 ppm of lead from PbCl_2_ or 1.3 ppm copper from CuSO_4_ or no metals as a control. When rehydrated with the water sources, with and without metals, each known metal-containing reaction produced a detectable signal within four hours, while the samples without metals showed no significant fluorescence activation (**Fig. 6c**,**d**). Unsurprisingly, in all cases, we observed lower signals from real-world water sources than the laboratory-grade water samples.

In addition to freeze-drying, field deployment also requires a convenient approach to detecting the fluorescence signals generated by ROSALIND. To this end, we developed a 3D-printed handheld device, designed to fit a strip of eight PCR tubes, that could visualize fluorescent output using the naked eye (**Supp. Fig. 9, Supp. Data**). The use of off-the-shelf components assembled onto a printed circuit board, including 470 nm LEDs and an adjustable potentiometer, allowed us to optimize the device for visualization of 3WJdB fluorescence using blue light for fluorescence-excitation and yellow stage light film as a longpass optical filter. Insertion of reaction tubes into the device leads to readily visible signals of the metal-containing reaction conditions (**Fig. 6c**,**d**).

Finally, we tested our freeze-dried ROSALIND copper sensor against ambient waters from Chile that are known to be highly enriched with copper from a combination of geogenic and anthropogenic processes [67]. Water samples were obtained from two streams: (1) the Yerba Loca Creek, upstream from Santiago, in the Maipo watershed of Central Chile, and (2) the Toro River, upstream from La Serena, in the Elqui watershed of Northern Chile (**Fig. 6e**). Copper concentrations were determined using Flame Atomic Absorption Spectroscopy (FAAS) (**Supp. Fig. 10**) and found to be in the range of 6.9-8.5 ppm, which were in the maximum activation region of the copper sensing dose response curve (**Fig. 6a**). In order to generate a series of tests that cover our detectable copper range, we serially diluted each sample (undiluted, 1:2, 1:4, 1:8, 1:16 and 1:32 dilutions) and measured copper by FAAS before using this series to rehydrate freeze-dried copper sensors (**Supp. Fig. 11**). When rehydrated with the 1:8 dilution, we observed clear visible signals (**Fig. 6e**), corresponding to copper concentrations of 0.88 ppm and 1.05 ppm from the Yerba Loca Creek and Toro River samples, respectively.

Overall, these results demonstrate that ROSALIND can be tuned for use with real-world water samples to detect contaminants of interest in formats that will allow for field-deployment.

## DISCUSSION

Here we report the development of a new synthetic biology biosensing platform, ROSALIND, for rapid and inexpensive detection of a wide range of chemical compounds and elements in aqueous samples. At the core of ROSALIND are simple and defined aTF-regulated IVT reactions—the combined use of aTFs that allosterically change DNA binding affinity to an operator sequence as a function of ligand concentration, and appropriately designed DNA templates that use this conditional binding to gate the production of a fluorescence-activating RNA output. This allowed us to modularly develop a set of 12 sensors simply by changing the aTF-operator pair. In fact, in each case, it was simply a matter of changing the operator sequence, with no other changes required to the DNA templates.

ROSALIND represents a significant advance over previous approaches for field-deployable chemical biosensing. In contrast to cellular biosensors, our platform does not require living cells to be maintained or contained, and does not suffer from transport or toxicity limitations of analytes. In addition, the use of bottom-up specified *in vitro* reactions allows each reaction component to be fine-tuned to achieve desired goals, which we utilized to shift the sensitivity of the copper and lead sensors (**Fig. 6a**,**b**). In contrast to other cell-free approaches, ROSALIND is more defined and much simpler. Specifically, the focus on transcriptional RNA outputs allowed us to avoid the complexity of translation, resulting in a simplified set of reaction components that can produce visible signals within tens of minutes. In addition, the defined nature of *in vitro* reactions removed much of the batch-to-batch variability that can occur when preparing cell extracts [68].

The ROSALIND platform relies on the production of a functional RNA that can bind and activate the fluorescence of an otherwise non-fluorescent dye. Although we used a variant of the Broccoli aptamer, other fluorescence-activating aptamers could be plugged into the ROSALIND platform [69]. Multiple RNA-based outputs could be used if their spectral properties and ligand-specificities are unique for multiplexing. In addition, the ROSALIND platform can be extended to many other aTFs that exist in nature, but are not yet characterized. The high-throughput and fast nature of ROSALIND and the ease of constructing reactions once an aTF is purified could offer a high-throughput screening format to uncover the ligands that activate these aTFs, to test for putative operator sequences, or to enable high-throughput aTF or promoter engineering efforts [70, 71]. As an example of this, we were able to use the ROSALIND platform to rapidly validate a putative aTF, CtcS, by showing its ability to bind to an identified operator site and induce with chlortetracycline. In addition, the platform should be easily extensible through further engineering of the aTF or operator sequences to enhance sensitivity, modify specificity, and otherwise tune system performance [36, 72–74].

A key limitation of the current ROSALIND platform is the use of aTFs that follow a simple repression mechanism involving ligand-mediated changes to aTF-DNA template binding. Although we were able to utilize an antisense binding approach using kleptamers [39] to expand the platform to be compatible with aTFs that bind the DNA template in the presence of a ligand, there remain numerous other aTFs that operate through a mechanism in which the allostery of ligand binding enables additional factors required for transcription to be recruited (i.e., s54) or even distorts the local topology of the DNA to prevent or enhance transcription initiation by the transcriptional machinery [38, 75].

A key feature of our study is demonstrating that freeze-dried ROSALIND reactions can function in the context of real-world water samples (**Fig. 6**). While reactions were found to function in all cases tested, we did observe reduced fluorescence signals when ROSALIND sensors were tested in non-laboratory water sources (i.e., drinking water, lake water and Chilean water). This can potentially be due to molecular components in the water samples that can (1) inhibit transcriptional activity or (2) quench fluorescence. In particular, when undiluted Chilean field samples were used to rehydrate the lyophilized copper sensor, we observed a relatively low fluorescent signal despite the high copper concentrations in the water samples (**Supp. Fig. 11**). We suspect that since a serial dilution was performed with laboratory-grade water on these field samples, the molecular components that interfere with ROSALIND are also diluted, enabling more robust signals to be generated in the diluted field samples. Nonetheless, our copper sensors were capable of detecting appropriate contaminants at the relevant concentration using a practical workflow of testing serial dilutions of the field samples. These matrix effects are an area of further exploration for future work.

Our work utilizes innovations in synthetic biology to fill a gap in existing water quality monitoring technology, which currently consists largely of sophisticated, centralized and expensive equipment to identify water contaminants. As a result, millions across the globe suffer from a lack of knowledge about their water quality, which can easily be compromised by contamination from environmental sources, anthropogenic activities, natural disasters or mismanagement. A technology such as ROSALIND, which allows the rapid and cheap identification of contaminated water at the source, will enable communities and individuals to monitor their water quality to meet their needs, and enable the gathering of critical information necessary for achieving the UN global sustainable development goal of safe and clean water.

## METHODS

### Strains and growth medium

*E. coli* strain K12 (NEB Turbo Competent *E. coli*, New England Biolabs #C2984) was used for routine cloning. *E. coli* strain Rosetta 2(DE3)pLysS (Novagen #71401) was used for recombinant protein expression. LB supplemented with the appropriate antibiotic(s) (100 µg/mL carbenicillin, 100 µg/mL kanamycin, and/or 34 µg/mL chloramphenicol) was used as the growth media.

### Plasmids and genetic parts assembly

DNA oligonucleotides for cloning and sequencing were synthesized by Integrated DNA Technologies. Genes encoding aTFs were synthesized either as gBlocks (Integrated DNA Technologies) or gene fragments (Twist Bioscience). Expression plasmids were cloned using Gibson Assembly (NEB Gibson Assembly Master Mix, New England Biolabs #E2611) into a pET-28c plasmid backbone, and were designed to overexpress recombinant proteins as either C-terminus or N-terminus His-tagged fusions. Certain constructs additionally incorporated a recognition sequence for cleavage and removal of the His-tag using TEV protease, and the construct for HucR expression contained a maltose binding protein sequence to enhance the solubility of the recombinant protein (**Supp. Table 2**). Gibson assembled constructs were transformed into NEB Turbo cells and isolated colonies were purified for plasmid DNA (QIAprep Spin Miniprep Kit, Qiagen #27106). Plasmid sequences were verified with Sanger DNA sequencing (Quintara Biosciences) using the primers listed in Supporting Data File 1.

Plasmids encoding transcription templates were constructed using pUC19-T7-3WJdB-T (Addgene plasmid #87308) as a backbone. These plasmids included a T7 RNAP promoter, an aTF operator site, the 3WJdB coding sequence [23], and a T7 terminator. Here we define the T7 RNAP promoter as a minimal 17 BP sequence (TAATACGACTCACTATA) excluding the first G that is transcribed. Plasmids were cloned using either Gibson Assembly or inverse PCR followed by blunt-end ligation, and then transformed into NEB Turbo cells for clonal isolation and sequence verification using the primer listed in Supporting Data File 1. Once sequence verified, transcription templates were generated by PCR amplification (Phusion High-Fidelity PCR Kit, New England Biolabs #E0553) of the plasmids to include a 5’ region upstream of the T7 promoter and a 3’ region ending with either 3WJdB or the T7 terminator with the primers listed in Supporting Data File 1. Amplified templates were then purified (QIAquick PCR purification kit, Qiagen #28106), verified for the presence of a single DNA band of expected size on a 2% TAE-Agarose gel, and concentrations were determined using the Qubit dsDNA BR Assay Kit (Invitrogen #Q32853).

The KB2 DNA transcription template was generated by denaturing two complementary oligonucleotides (Supporting Data File 1) at 95° C for 3 minutes and slow cooling (−0.1° C/s) to room temperature in 1X annealing buffer (100 mM potassium acetate and 30 mM HEPES, pH 8.0). Annealed oligonucleotides where then purified by resolving them on a 20% native PAGE-TBE gels, isolating the band of expected size, and eluting at room temperature overnight in 1X annealing buffer. The eluted DNA template was then ethanol precipitated, resuspended in MilliQ H_2_O, and quantified using the Qubit dsDNA BR Assay Kit.

A table listing the sequences and Addgene accession numbers of all plasmids generated in this study are listed in Supporting Data File 1.

### aTF expression and purification

Supplementary Table 2 provides a listing of the exact expression and purification process for each aTF. In general, with the exception of CadC and AdcR, each protein was expressed and purified as follows. Sequence-verified pET-28c plasmids were transformed into the Rosetta 2(DE3)pLysS *E. coli* strain for protein expression. A single transformed colony from each expression plasmid transformation was then used to seed a small overnight culture grown at 37° C. Overnight cultures were then seeded into larger expression cultures (10 mL starter culture / 1 L expression culture). Cultures were induced for overexpression at an optical density (600 nm) of ∼0.5 with 100 to 500 µM of IPTG and grown for 4 additional hours at 37° C or overnight at 30° C. Cultures were then pelleted by centrifuging at 4,000 RCF for 20 minutes at 4 °C in a Thermo Scientific Sorvall Lynx 4000, and then either were immediately stored at −80° C or resuspended in lysis buffer (10 mM Tris-HCl pH 7.5 - 8.5, 500 mM NaCl, 1 mM TCEP, and protease inhibitor (cOmplete EDTA-free Protease Inhibitor Cocktail, Roche)). Resuspended cells were then lysed on ice through four rounds of ultrasonication (50% duty cycle) for 1 min, with 2 min of rest on ice in between rounds. Lysates were then centrifuged at 13,000 RCF for 30 minutes to remove insoluble material. Clarified supernatants were then purified using His-tag affinity chromatography with a Ni-NTA column (HisTrap FF 5mL column, GE Healthcare Life Sciences) followed by size exclusion chromatography (Superdex HiLoad 26/600 200 pg column, GE Healthcare Life Sciences) using an AKTAxpress fast protein liquid chromatography (FPLC) system. The collected fractions from the FPLC were concentrated and buffer exchanged (25 mM Tris-HCl, 100 mM NaCl, 1mM TCEP, 50% glycerol v/v) using centrifugal filtration (Amicon Ultra-0.5, Millipore Sigma). Protein concentrations were determined using a Qubit Protein Assay Kit (Invitrogen #Q33212). The purity and size of the proteins were validated on SDS-PAGE gel (Mini-PROTEAN TGX and Mini-TETRA cell, Bio-Rad). Purified proteins were stored at –20° C. CadC and AdcR, whose genes were cloned into a pET-3d and pET-3a plasmid backbone respectively, were purified with the protocol previously described [76]. For CadC purification, the protocol was modified to have a higher concentration of TCEP (5 mM) in its buffers.

### In vitro transcription (IVT) reactions

NEB HiScribe T7 Quick High Yield RNA Synthesis Kit (#E2050S, New England Biolabs) was used for data shown in Figure 1 according to the manufacturers protocol with the addition of 2.25 mM DFHBI-1T (Tocris #5610) in the reaction mix. Homemade IVT reactions were set up by adding the following components listed at their final concentration: IVT buffer (40 mM Tris-HCL pH 8, 8 mM MgCl_2_, 10 mM DTT, and 20 mM NaCl), 2.25 mM DFHBI-1T, 11.4 mM Tris-buffered NTPs pH 7.5, 0.3U thermostable inorganic pyrophosphatase (#M0296S, New England Biolabs), DNA transcription template(s), and MilliQ H_2_O to a total volume of 20 µL. Regulated IVT reactions additionally included a purified aTF at the indicated concentration and were equilibrated at 37° C for 15 minutes. Immediately prior to plate reader measurements, 0.2 ng of T7 RNA polymerase and, optionally, a ligand at the indicated concentration (**Supp. Table 3**) were added to the reaction. Reactions were then loaded onto a 384-well, black, optically-clear, flat-bottom plate and measured on a plate reader (Synergy H1, BioTek) at 3 min intervals with excitation and emission wavelengths of 472 nm and 507 nm, respectively. Supporting Data File 2 contains a master experimental planning sheet that describes how a typical ROSALIND reaction is setup.

### Freeze-drying

Lyophilization of ROSALIND reactions was performed by assembling the components of regulated *in vitro* transcription (see above) with the addition of 50 mM sucrose and 200 mM mannitol. Assembled reaction tubes were immediately transferred into a pre-chilled aluminum block and placed in a −80°C freezer for 10 minutes to allow slow-freezing. Following the slow-freezing, reaction tubes were submerged in liquid nitrogen and then transferred to a FreeZone 2.5 L Bench Top Freeze Dry System (Labconco) for overnight freeze-drying with a condenser temperature of −85° C and 0.04 millibar pressure. Unless rehydrated immediately, freeze-dried reactions were stored in glass jars filled with desiccant (Drierite) at room temperature.

### Mean equivalent fluorescence (MEF) standardization

To convert arbitrary fluorescence units to mean equivalent fluorescence (MEF) of fluorescein isothiocyanate (FITC), we used a NIST traceable standard (Invitrogen #F36915). Serial dilutions from a 50 µM stock were prepared in 100 mM sodium borate buffer at pH 9.5, including a 100 mM sodium borate buffer blank (total of 12 samples). The samples were prepared in technical and experimental triplicate (12 samples X 9 replicates = 108 samples total), and fluorescence values were read at an excitation wavelength of 472 nm and emission wavelength of 507 nm for 3WJdB-activated fluorescence, or 485 nm excitation and 515 nm emission for sfGFP fluorescence. Fluorescence values for a concentration in which a single replicate saturated the plate reader were excluded from analysis. The remaining replicates (9 per sample) were then averaged at each FITC concentration, and the average fluorescence value of the blank was subtracted from all values. Linear regression was then performed for concentrations within the linear range of fluorescence (0–6.25 µM FITC) between the measured fluorescence values in arbitrary units and the concentration of FITC to identify the conversion factor. For each plate reader, excitation, emission, and gain setting, we found a linear conversion factor that was used to correlate arbitrary fluorescence values to MEF (µM FITC). We note that no background subtraction was performed when analyzing outputs from any ROSALIND reaction. An example of this standardization procedure is shown in Supplementary Figure 1.

### Chilean Water Sampling

River water samples were taken from the Yerba Loca Creek in the upper Mapocho-Maipo watershed, and the Toro River in the upper Elqui River watersheds. Both streams are metal-polluted tributaries to rivers that provide ecosystem services (*e.g.*, drinking water sources, habitat for ecosystems, recreation) to major Chilean cities. Acid-washed plastic bottles were used to collect 200 mL of filtered (0.45 µm nylon membranes) samples and shipped in a cooler. Given the low pH (2.9 < pH < 4) and the objective of the experiments, pH was not adjusted.

### Flame atomic absorption spectroscopy measurement

1:1 serial dilution with laboratory-grade water was performed on Chilean water samples (4 dilutions per location: 1:2, 1:4, 1:8, 1:16). Flame atomic absorption spectroscopy (PerkinElmer PinAAcle 500) was calibrated for Cu(II) using CuNO_3_ reference solution as a standard (Fisher Scientific, #AC207681000) with the maximum threshold value of 2 ppm. Chilean water samples were measured starting with the 1:16 dilutions and, based on the estimated values of the 1:16 dilutions, the 1:2 and lower dilutions were excluded from the measurements to avoid oversaturating the instrument. Each dilution was measured three times and averaged, and linear regressions were performed on these averaged values to calculate the concentrations of the undiluted samples (**Supp. Fig. 10**).

### Photographs of reactions

Photographs of reaction tubes in Figures 1 and 6 were taken on an Apple iPhone X or 6S using the default Apple iOS Camera application. The resulting images were only cropped. Source image files are available as Supplementary Data File 3.

## ACKNOWLEDGEMENTS

We would like to thank Professor Jean-François Gaillard and Zhaoxun Yang (Northwestern) for assistance with flame atomic absorbance spectroscopy measurements. We also thank Nina Donghia (MIT) for helpful discussions on lyophilization of cell free reactions, Sergii Pshenychny (Recombinant Protein Production Core at Northwestern University) for assistance in protein purification, and John Bussan, Frank Lantz and Bob Golenia (Northwestern University Research Shop) for assistance in the handheld illuminator development. J. K. J. was supported in part by the Northwestern University Graduate School Cluster in Biotechnology, System, and Synthetic Biology, which is affiliated with the Biotechnology Training Program. This work was supported by funding from the Pew Charitable Trusts (to D.A.C.), NIH R35 GM118157 (to D.P.G.), CONICYT-FONDECYT (1161337 to P. A. P.), CONICYT-FONDAP (15110020 to P. A. P.), an NSF CAREER award (1452441 to J.B.L.), and Searle Funds at The Chicago Community Trust (to J.B.L.).

## AUTHOR CONTRIBUTIONS

Conceptualization, K.K.A., J.K.J., J.J.C & J.B.L.; Data curation, K.K.A., J.K.J., M.S.V. & J.B.L.; Formal analysis, K.K.A., J.K.J. & J.B.L.; Funding acquisition, J.B.L., P. A. P.; Investigation, K.K.A., J.K.J., M.S.V., P.R.C., J.W.L.; Methodology, K.K.A., J.K.J., M.S.V., J.W.L., D.A.C. & J.B.L.; Project administration, K.K.A., J.K.J. & J.B.L.; Resources, D.A.C., D.P.G., P.A.P.; Supervision, K.K.A, J.J.C. & J.B.L; Validation, K.K.A., J.K.J., M.S.V., P.R.C.; Visualization, K.K.A., J.K.J. & J.B.L.; Writing – original draft, K.K.A., J.K.J. & J.B.L.; Writing – review & editing, K.K.A., J.K.J., M.S.V., P.R.C., J.W.L., D.A.C., P.A.P., D.P.G., J.J.C. & J.B.L. K.K.A. and J.K.J. contributed equally to this work.

## COMPETING INTERESTS STATEMENT

K.K.A., J.K.J. & J.B.L. have submitted a US provisional patent application (No. 62/758,242) relating to regulated in vitro transcription reactions. K.K.A., J.K.J., M.S.V., P.R.C., J.W.L., J.J.C. & J.B.L. have submitted a US provisional patent application (No. pending) relating to the preservation and stabilization of in vitro transcription reactions.

## Notes

#### Summary of Updates

Updated acknowledgements and author ORCID.

