## Supplementary Information for "Rapid, Low-Cost Detection of Water Contaminants Using Regulated *In Vitro* Transcription"

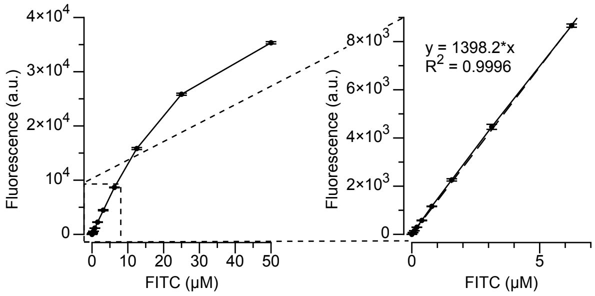


**Supp. Fig. 1 | Mean Equivalent Fluorescence (MEF) standardization.** Arbitrary units of fluorescence were standardized to µM concentrations of fluorescein isothiocyanate (FITC) using a NIST traceable standard (see Methods). In the representative example shown here, a dilution series of FITC was prepared in buffer (100 mM sodium borate, pH 9.5) and measured on a plate reader using the same settings for measuring 3WJdB signal (472 nm excitation, 507 nm emission). The resulting curve, calculated over the linear range 0-6.25 µM, was then used to standardize fluorescence measured from ROSALIND reactions. The standard curve was generated at regular intervals for each plate reader and each measurement setting. Data shown are for 3 experimental replicates with 3 technical replicates each (n = 9). Error bars indicate standard deviation.

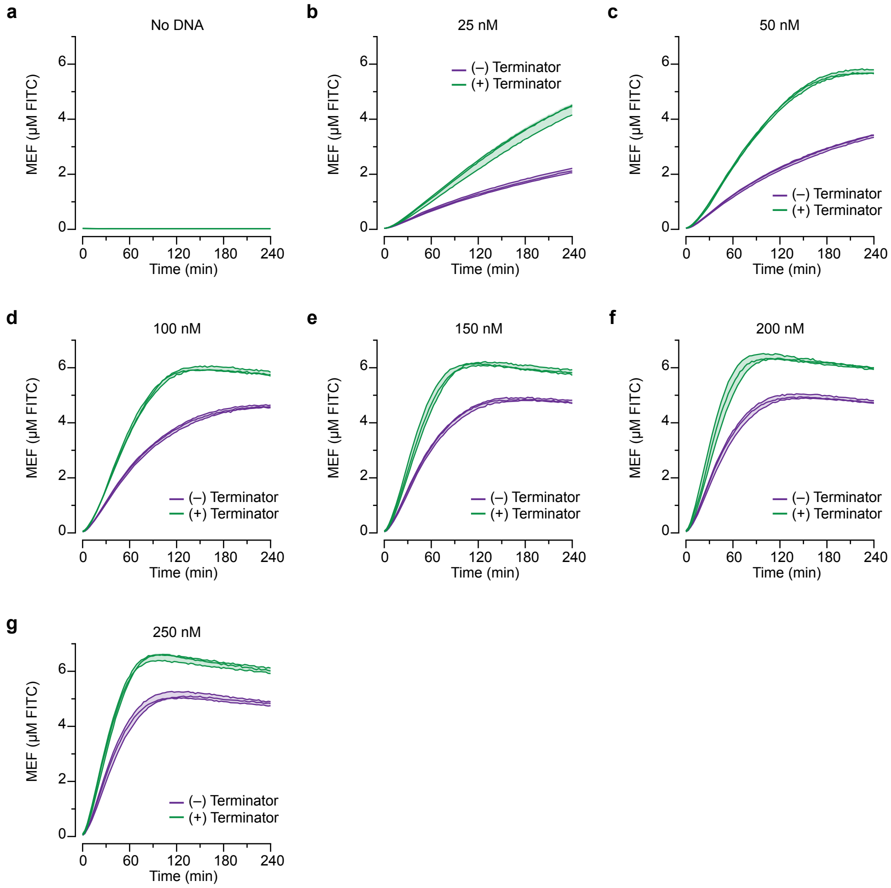


**Supp. Fig. 2 | Fluorescence-activation of 3WJdB during run-off *in vitro* transcription reactions is improved by additionally encoding a T7 terminator.** *In vitro* transcription reactions using a commercially available kit (NEB HiScribe T7 Quick High Yield RNA Synthesis Kit) were performed across a range of DNA transcription template concentrations, with and without an additional T7 terminator sequence. **a,** Reactions containing no DNA failed to generate signal, while all other DNA containing reactions (**b-g**) generated measurable fluorescence that improved with the addition of a template-encoded T7 terminator. We hypothesize that the terminator-dependent increase in fluorescence could be due to the presence of the terminator structure encouraging faster recycling of T7 RNAP from the elongation phase, to the termination phase, and then to initiation phase. Although less likely due to the highly-structured design of 3WJdB, the terminator-dependent increase could be due to the additional terminator structure stabilizing the productive, fluorescence-activating fold of 3WJdB. All data shown for 3 experimental replicates as lines with raw fluorescence values calibrated to µM FITC. Shading indicates the average value ± standard deviation.

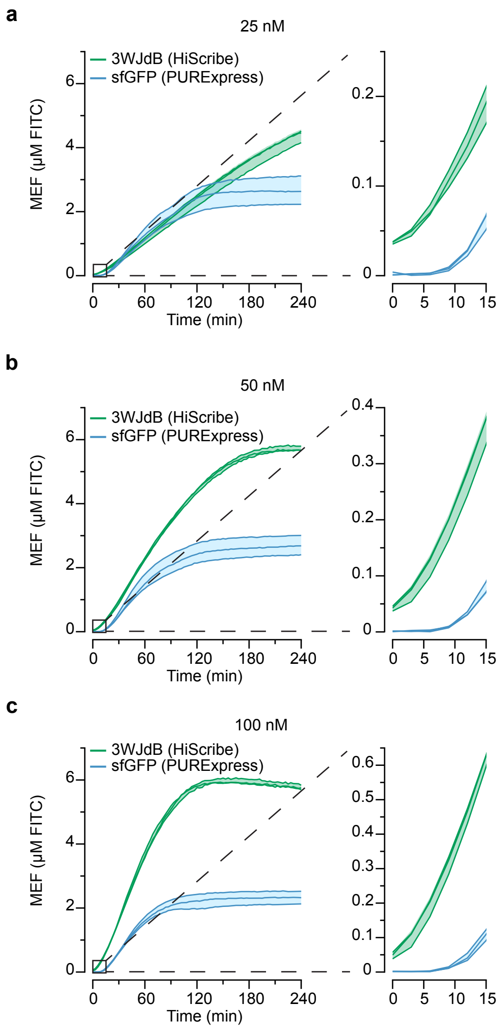


**Supp. Fig. 3 | Fluorescence-activation from 3WJdB transcription reactions is faster and brighter than fluorescence from transcription-translation (PURE) reactions of sfGFP.** All data shown for 3 experimental replicates as lines with raw fluorescence values calibrated to μM FITC. Shading indicates the average value ± standard deviation.

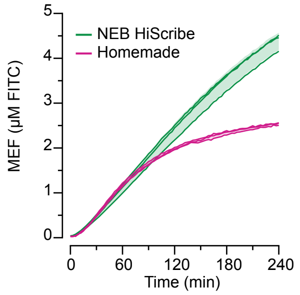


**Supp. Fig. 4 | Homemade transcription reactions can be optimized to perform as well as commercial kits.** Homemade *in vitro* transcription reactions were compared to a commercially available high yield transcription kit (NEB HiScribe T7 Quick High Yield RNA Synthesis Kit). 0.5 pmol of DNA encoding T7-3WJdB-T was added to each reaction in a total reaction volume of 20 μL (25 nM transcription template). Over the course of 1 hour the data show similar fluorescence activation. However, the homemade reaction begins to saturate after 1 hour, likely due to the exhaustion of NTPs in the homemade reaction (11.4 mM) when compared to the commercial kit (40 mM). All data shown for 3 experimental replicates with raw fluorescence values calibrated to μM FITC. Shading indicates the average value ± standard deviation.

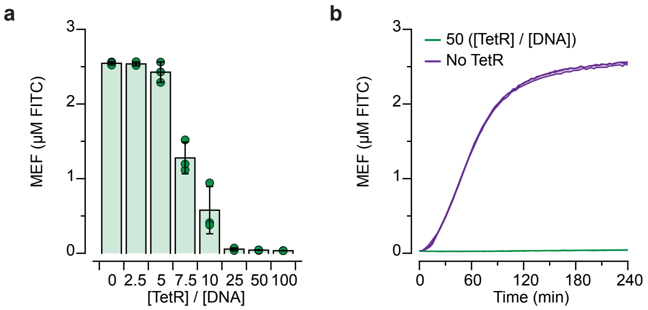


**Supp. Fig. 5 | Titration of the TetR aTF into a regulated *in vitro* transcription.** TetR protein was titrated as a function of DNA concentration to optimize for low fluorescence-activation. **a**, Data taken 4 hours after initiating reactions with T7 RNAP. Fluorescence-activation is substantially repressed at 25-fold excess or greater of TetR dimer over DNA template. **b**, Time course comparing the 0-fold and 50-fold [TetR]/[DNA] ratio experiments. All data shown for 3 experimental replicates as points or lines with raw fluorescence values calibrated to µM FITC. Colored bars and shaded regions indicate the average value ± standard deviation. All data shown for 3 experimental replicates with raw fluorescence values calibrated to µM FITC.

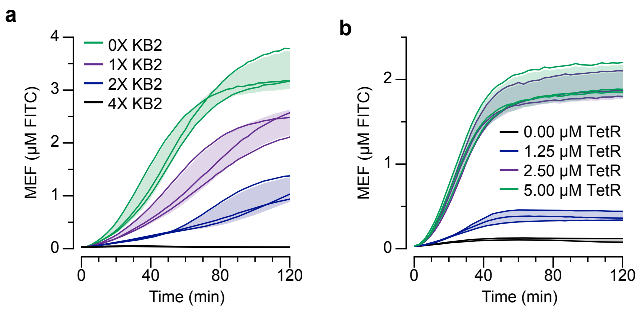


**Supp. Fig. 6 | RNA kleptamers are used to invert transcription factor responses. a,** Titration of the KB2 DNA template into an *in vitro* transcription of 3WJdB decreases the fluorescence from 3WJdB and **b,** Titration of TetR protein into an *in vitro* transcription of 3WJdB and a TetR regulated KB2 expression template restores the fluorescence from 3WJdB.

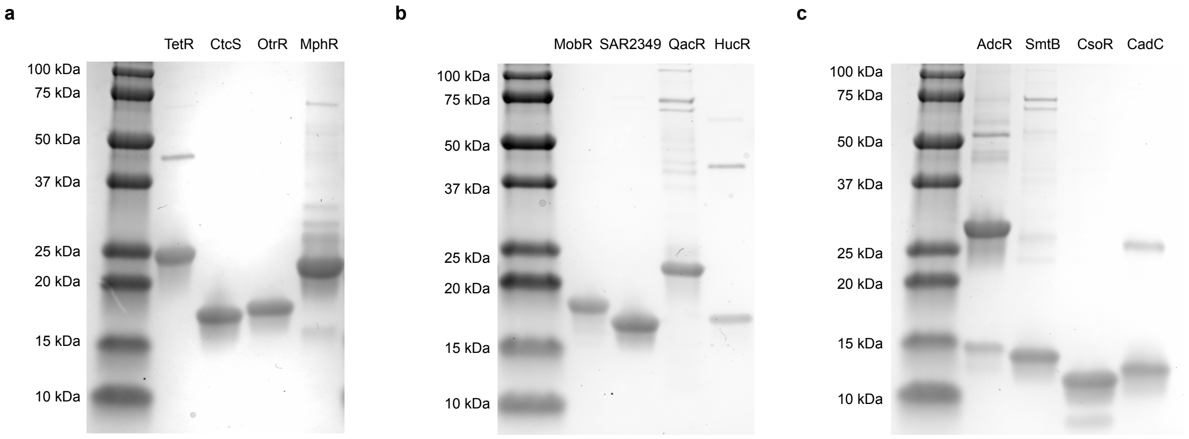


**Supp. Fig. 7 | SDS-PAGE protein gel of all aTFs. a**, antibiotics sensing aTFs (TetR – 24.9 kDa, CtcS – 18.7 kDa, OtrR – 19.5 kDa, MphR – 23.2 kDa), **b,** small molecule sensing aTFs (MobR – 20.6 kDa, SAR2349 – 18.9 kDa, QacR – 23.8 kDa, HucR – 19.8 kDa), and **c,** metal sensing aTFs (AdcR – 16.6 kDa, SmtB – 15.2 kDa, CsoR – 11.5 kDa, and CadC – 13.8 kDa). The molecular weights listed are calculated from aTFs amino acid sequences (**Supp. Data File 1**). The gel was run at a denaturing and non-reducing condition. In the AdcR lane, the most intense band corresponds to AdcR dimer (~33.2 kDa) formed by a disulfide bond under this condition.

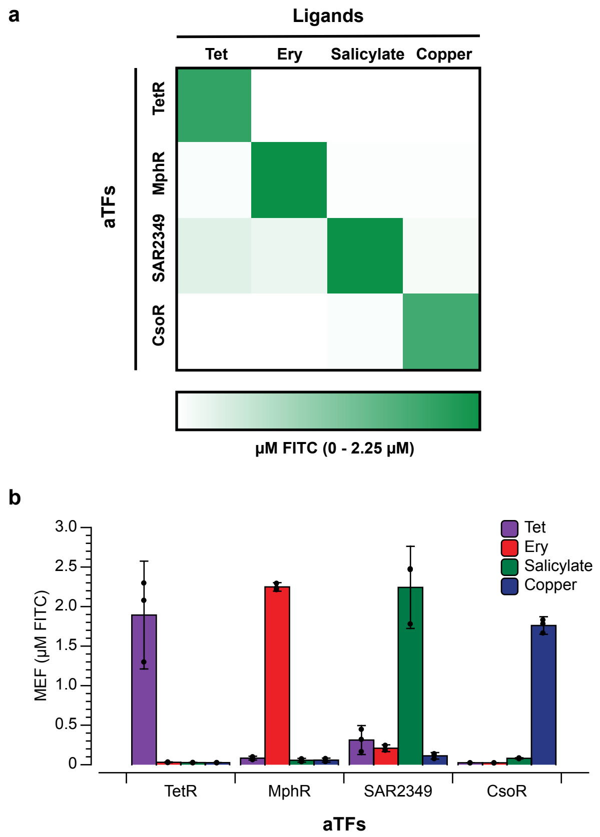


**Supp. Fig. 8 | Orthogonality of ROSALIND across different families of ligands.** Four different sensors (one representative from each family) were tested for crosstalk with different ligands and determined to be orthogonal with each other as shown in the **a**, 4X4 orthogonality matrix and **b**, corresponding bar graph. Concentrations of ligands tested are 25 µM of tetracycline, 50 µM of erythromycin, 12 mM of salicylate, and 10 µM of $CuSO_4$. Reactions were configured as described in Figure 5.

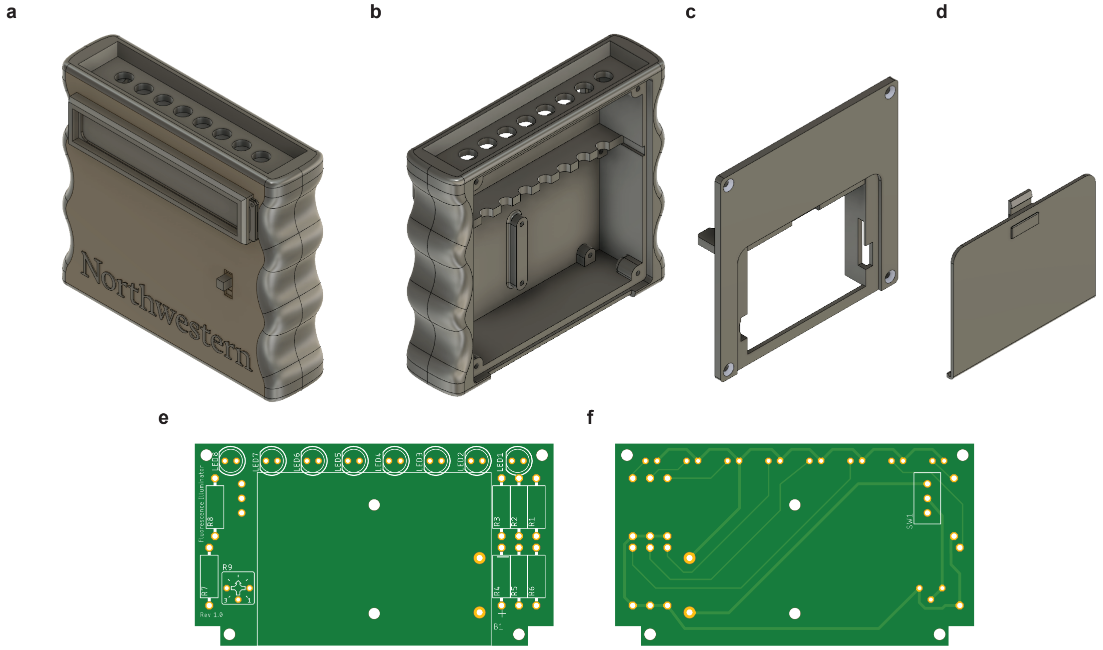


**Supp. Fig. 9 | Handheld fluorescence illuminator design. a,** A low-cost handheld fluorescence illuminator enables point-of-use functionality for ROSALIND. 3D printed components include a **b,** front case, **c,** back case, and **d,** battery cover. A printed circuit board (**e,** front view facing back case, **f,** rear view facing front case) mounts in the front case and connects LEDs, resistors, a trimmer potentiometer, a power switch, and a battery holder. Full CAD files, 3D print files (.STL), PCB design files, and assembly instructions are provided in the **Supporting Data File 4**.

a

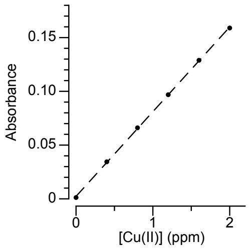


b

| Dilution | Yerba Loca (ppm) | Toro (ppm) |
|---|---|---|
| *1:4* | 1.73(7) ± 0.010 | 2.03(3) ± 0.010 |
| *1:8* | 0.87(9) ± 0.010 | 1.04(5) ± 0.010 |
| *1:16* | 0.43(4) ± 0.010 | 0.51(8) ± 0.010 |
| *Undiluted** | 6.96(4) | 8.43(7) |

**These values were calculated by performing linear regression (see below)*

c

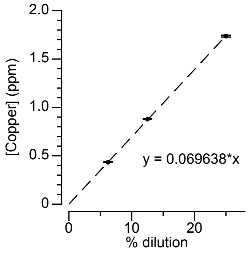


d

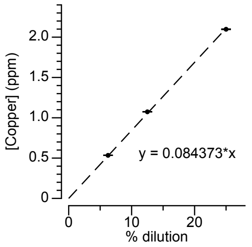


**Supp. Fig. 10 | Flame atomic absorption spectroscopy (FAAS) measurements of Chilean field samples.** FAAS was calibrated with a $Cu(NO_3)_2$ standard solution as shown in **a**. Serial dilution was performed on each sample to create three separate diluted samples that are in the operating ppm range of FAAS, and their ppm values were measured as reported in Table **b**. Using these measurements, linear regression on the averages of each dilution was performed to calculate the ppm value of the undiluted sample from **c**, the Yerba Loca Creek and **d**, the Toro River.

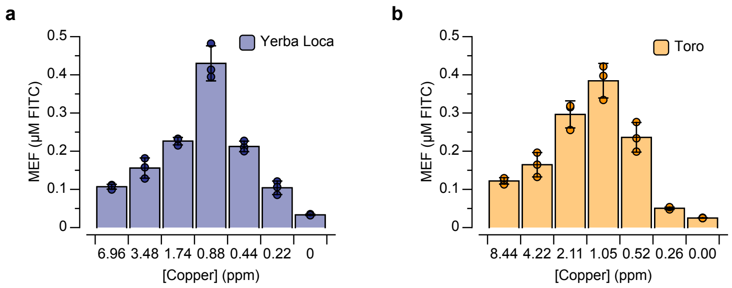


**Supp. Fig. 11 | Titrations of Chilean field samples into lyophilized ROSALIND copper sensing reactions.** 1:1 serial dilution was performed on the field samples from **a**, the Yerba Loca Creek and **b**, the Toro River. The diluted field samples and laboratory-grade water (0.0 ppm) were used to rehydrate lyophilized ROSALIND copper sensing reactions. 4-hour end-point fluorescent values are reported over three replicates (points) calibrated to µM FITC. Bars represent averages over the replicates and error bars standard deviations. The copper concentrations indicated were taken from Supp. Fig. 10, or calculated from the extrapolated ppm value of the undiluted sample.

### Supplementary Table 1 – Estimated cost per ROSALIND reaction[*]

| Component | Supplier | Catalog Number | Quantity | Price (USD) | [Stock] | Stock Volume (mL) | Amount / Reaction (mL) | # Reactions | Cost / 20 µL Reaction (USD) |
|---|---|---|---|---|---|---|---|---|---|
| DFHBI-1T | Tocris | 5610 | 10 mg | $240.59 | 40 mM | 0.781 | 0.00113 | 691 | $0.35 |
| ATP | Sigma | A2383-25G | 25 g | $466.48 | 100 mM | 453.605 | 0.00057 | 795798 | < $0.01 |
| CTP | Sigma | C1506-1G | 1 g | $448.84 | 100 mM | 18.971 | 0.00057 | 33282 | $0.01 |
| GTP | Sigma | G8877-1G | 1 g | $600.74 | 100 mM | 19.114 | 0.00057 | 33533 | $0.02 |
| UTP | Sigma | U6625-1G | 1 g | $495.88 | 100 mM | 18.179 | 0.00057 | 31893 | $0.01 |
| Tris | Sigma | RDD009-2.5KG | 2.5 kg | $166.56 | 2 M | 7931.5 | 0.00020 | 39657500 | << $0.01 |
| $MgCl_2$ | Sigma | M2670-1KG | 1 kg | $109.81 | 800 mM | 6148.5 | 0.00020 | 30742500 | << $0.01 |
| Spermidine | Sigma | S2626-25G | 25 g | $498.82 | 200 mM | 860.585 | 0.00020 | 4302926 | << $0.01 |
| NaCl | Sigma | 746398-25KG | 25 kg | $198.50 | 2 M | 213894.6 | 0.00020 | 1069473000 | << $0.01 |
| DTT | Gold Bio | DTT500 | 500 g | $1,425 | 1 M | 3241.5 | 0.00020 | 16207500 | << $0.01 |
| Sucrose | Sigma | S0389-5KG | 5 kg | $61.53 | 5 M | 2921.4 | 0.00020 | 14607000 | << $0.01 |
| Mannitol | Sigma | M4125-5KG | 5 kg | $173.01 | 1 M | 27446.9 | 0.00500 | 5489380 | << $0.01 |
| TIPP | NEB | M0296L | 1250 U | $244.79 | 2 U/µL | 0.625 | 0.00015 | 4167 | $0.06 |
| T7 RNAP[a] | n/a | n/a | 50 mg | $400.00 | 1 mg/mL | 50 | 0.0002 | 250000 | < $0.01 |
| aTF[a] | n/a | n/a | 10 mg | $400.00 | 1 mg/mL | 10 | 0.0002 | 50000 | < $0.01 |
| DNA Template[b] | n/a | n/a | 20 µg | $42.81 | 0.1 µg/µL | 0.2 | 0.001 | 200 | $0.21 |
| **TOTAL** | | | | | | | | | **~ $0.67 / reaction** |

[a]Homemade. Estimated for 2 L expression culture using core facility pricing (NU rPPC, March 2019).
[b]Calculated for the cost of a 2 mL PCR and purification using NEB #M3050 ($34.96/prep), NEB #N0447L ($2.45/prep), Qiagen #28106 ($4.40/prep), and PCR primers (< $1/prep).
*Calculated using institutional pricing (March 2019). Not included in calculation: labor, overhead, equipment, consumables (*e.g.* pipette tips) and additional materials (*e.g.* for pH adjustments), laboratory grade water, storage, shipping.

**Supp. Table 2 | Plasmid constructs and purification method of aTFs.**

| aTF | Type of Purification | Tag Location | TEV Cleavage | Columns Used |
|---|---|---|---|---|
| TetR | His-tag affinity followed by size exclusion | C-terminus 6XHis-tag | No | HisTrap FF 5mL column for affinity, and Superdex HiLoad 26/600 200 pg column for size exclusion |
| OtrR | His-tag affinity followed by size exclusion | C-terminus TEV followed by 6XHis-tag | No | HisTrap FF 5mL column for affinity, and Superdex HiLoad 26/600 200 pg column for size exclusion |
| CtcR | His-tag affinity followed by size exclusion | C-terminus 6XHis-tag | No | HisTrap FF 5mL column for affinity, and Superdex HiLoad 26/600 200 pg column for size exclusion |
| MphR | His-tag affinity followed by size exclusion | C-terminus 6XHis-tag | No | HisTrap FF 5mL column for affinity, and Superdex HiLoad 26/600 200 pg column for size exclusion |
| MobR | His-tag affinity followed by size exclusion | C-terminus TEV followed by 6XHis-tag | No | HisTrap FF 5mL column for affinity, and Superdex HiLoad 26/600 200 pg column for size exclusion |
| SAR2349 | His-tag affinity | C-terminus TEV followed by 6XHis-tag | No | Gravity flow column packed with Qiagen Ni-NTA Agarose |
| QacR | His-tag affinity followed by size exclusion | C-terminus 6XHis-tag | No | HisTrap FF 5mL column for affinity, and Superdex HiLoad 26/600 200 pg column for size exclusion |
| HucR | His-tag affinity | N-terminus 6XHis-tag and MBP followed by TEV tag | Yes | Gravity flow column packed with Qiagen Ni-NTA Agarose |
| AdcR | PEI precipitation, $(NH_4)_2SO_4$ precipitation, ion exchange followed by size exclusion | N/A | N/A | SP-Sepharose fast flow column for ion exchange, and Superdex 75 preparative-grade column for size exclusion |
| SmtB | His-tag affinity | C-terminus TEV followed by 6XHis-tag | No | Gravity flow column packed with Qiagen Ni-NTA Agarose |

| | | | | |
|---|---|---|---|---|
| CsoR | His-tag affinity | N-terminus 6XHis-tag followed by TEV tag | Yes | Gravity flow column packed with Qiagen Ni-NTA Agarose |
| CadC | PEI precipitation, $(NH_4)_2SO_4$ precipitation, ion exchange followed by size exclusion | N/A | N/A | SP-Sepharose fast flow column for ion exchange, and Superdex 75 preparative-grade column for size exclusion |

**Supp. Table 3 | Concentrations of ROSALIND components in each sensor.**

| Ligand – aTF Pair | [DNA] (nM) | [aTF] (μM) | [Ligand] (μM) | Ligand Solvent | [Ligand Stock] (μM) | Ligand Purchased From |
|---|---|---|---|---|---|---|
| Anhydrotetracycline – TetR | 25 | 1.25 | 0.1-50 | Ethanol | 5 – 500 | Sigma-Aldrich (Cat#: 37919 – 100MG – R) |
| Tetracycline – TetR | 25 | 1.25 | 25 | Laboratory-grade $H_2O$ | 46.2 | Gold Biotechnology (Cat#: T – 101 – 25) |
| Doxycycline – TetR | 25 | 1.25 | 12.5 | Laboratory-grade $H_2O$ | 45.8 | Gold Biotechnology (Cat#: D – 500 – 1) |
| Oxytetracycline – OtrR | 25 | 2.5 | 100 | Laboratory-grade $H_2O$ | 159 | Gold Biotechnology (Cat#: O – 410 – 10) |
| Chlortetracycline – CtcS | 25 | 1.25 | 50 | Laboratory-grade $H_2O$ | 500 | Gold Biotechnology (Cat#: C – 840 – 5) |
| Erythromycin – MphR | 25 | 0.625 | 50 | Ethanol | 68.126 | Sigma-Aldrich (Cat#: E5389 – 1G) |
| Azithromycin – MphR | 25 | 0.625 | 50 | Ethanol | 500 | Sigma-Aldrich (Cat#: PHR1088 – 1G) |
| Clarithromycin – MphR | 25 | 0.625 | 50 | Ethanol | 500 | Sigma-Aldrich (Cat#: PHR1038 – 500MG) |
| Roxithromycin – MphR | 25 | 0.625 | 250 | Ethanol | 3488.5 | Sigma-Aldrich (Cat#: R4393-1G) |
| 3-Hydroxy Benzoic Acid – MobR | 10 | 100 | 2,000 | Tris-base Buffer | 13,153 | Sigma-Aldrich (Cat#: H20008 – 5G) |
| Salicylate – SAR2349 | 10 | 50 | 12,000 | Laboratory-grade $H_2O$ | 236,102 | Sigma-Aldrich (Cat#: S2679 – 100G) |
| Benzalkonium Chloride – QacR | 25 | 5 | 100 | Laboratory-grade $H_2O$ | 387.5 | Sigma-Aldrich (Cat#: 12060 – 5G) |
| Uric Acid – HucR | 17.5 | 2.15 | 303 | 30mM NaOH | 1011 | Sigma-Aldrich (Cat#: U0881 – 10G) |
| $ZnSO_4$ – AdcR | 7.5 | 1.5 | 30 | Laboratory-grade $H_2O$ | 200 | Sigma-Aldrich (Cat#: 83265 – 250mL – F) |
| $ZnSO_4$ – SmtB | 25 | 5 | 10 | Laboratory-grade $H_2O$ | 202 | Sigma-Aldrich (Cat#: 83265 – 250mL – F) |
| $CuSO_4$ – CsoR | 25 | 2.5 | 10 | Laboratory-grade $H_2O$ | 0.1 – 1,000 | Sigma-Aldrich (Cat#: 209198 – 5G) |

| | | | | | | |
|---|---|---|---|---|---|---|
| $CdCl_2$ – CadC | 20 | 1.5 | 10 | Laboratory-grade $H_2O$ | 360 | Sigma-Aldrich (Cat#: 202908 – 10G) |
| $PbCl_2$ – CadC | 10 | 1.5 | 51.8 | Laboratory-grade $H_2O$ | 0.2 – 200 | Sigma-Aldrich (Cat#: 203572 – 10G) |

**Supporting Data File Descriptions**

Alam_Jung_ROSALIND_SuppVideo_4kUHD.mp4 – Supplementary Video showing illumination of an unregulated ROSALIND reaction.

Alam_Jung_ROSALIND_Supp_Data_File1_Sequences.xlsx – DNA and Protein sequences used in this study.

Alam_Jung_ROSALIND_Supp_Data_File2_Experiment_Mastersheet.xlsx – Template excel worksheet for setting up a ROSALIND reaction.

Alam_Jung_ROSALIND_Supp_Data_File3_Images.zip – Raw illuminator and SDS-PAGE gel images included in figures.

Alam_Jung_ROSALIND_Supp_Data_File4_Handheld_Fluorescence_Illuminator_Instructions.zip – Instructions for 3D printing and assembling a hand-held illuminator.

Alam_Jung_ROSALIND_Supp_Data_File5_All_Source_Data.xlsx – Calibrated plate reader data for all figures.
