## Supporting Data Files for "Rapid, Low-Cost Detection of Water Contaminants Using Regulated *In Vitro* Transcription": IMG_5546_NOTES.txt

"Rapid, Low-Cost Detection of Water Contaminants using Regulated In Vitro Transcriptions"
Alam, Jung, et al. (2019)

IMG_5546.JPG show the visible fluorescence of 3WJdB in vitro transcription reactions.

From left to right:
1. Blank (Dye + Buffer only): MEF = 0 µM FITC
2. MEF = 0.125 µM FITC
3. MEF = 0.25 µM FITC
4. MEF = 0.5 µM FITC
5. MEF = 1 µM FITC
6. MEF = 2 µM FITC
7. MEF = 3 µM FITC
8. MEF = 6 µM FITC
