## Supporting Data Files for "Rapid, Low-Cost Detection of Water Contaminants Using Regulated *In Vitro* Transcription": IMG_1360_NOTES.txt

"Rapid, Low-Cost Detection of Water Contaminants using Regulated In Vitro Transcriptions"
Alam, Jung, et al. (2019)

IMG_1360.JPG show the visible fluorescence signals from the copper sensors that have been rehydrated with three different water sources.

From left to right, copper sensor rehydrated with:
1. Laboratory-grade water
2. Laboratory-grade water spiked with 1.3ppm copper
3. Drinking water
4. Drinking water spiked with 1.3ppm copper
5. Lake water
6. Lake water spiked with 1.3ppm copper
