## Supporting Data Files for "Rapid, Low-Cost Detection of Water Contaminants Using Regulated *In Vitro* Transcription": IMG_1425_NOTES.txt

"Rapid, Low-Cost Detection of Water Contaminants using Regulated In Vitro Transcriptions"
Alam, Jung, et al. (2019)

IMG_1425.JPG show the visible fluorescence signals from the lead sensors that have been rehydrated with three different water sources.

From left to right, lead sensor rehydrated with:
1. Laboratory-grade water
2. Laboratory-grade water spiked with 2ppm lead
3. Drinking water
4. Drinking water spiked with 2ppm lead
5. Lake water
6. Lake water spiked with 2ppm lead
