## Supporting Data Files for "Rapid, Low-Cost Detection of Water Contaminants Using Regulated *In Vitro* Transcription": IMG_1364_NOTES.txt

"Rapid, Low-Cost Detection of Water Contaminants using Regulated In Vitro Transcriptions"
Alam, Jung, et al. (2019)

IMG_1364.JPG show the visible fluorescence signals from the copper sensors that have been rehydrated with the water sample from the Yerba Loca Creek, Chile.

From left to right, copper sensor rehydrated with:
1. Undiluted (6.96 ppm)
2. 1:2 dilution (3.48 ppm)
3. 1:4 dilution (1.74 ppm)
4. 1:8 dilution (0.88 ppm)
5. 1:16 dilution (0.44 ppm)
6. 1:32 dilution (0.22 ppm)
7. Laboratory-grade water (0 ppm)
