## Supporting Data Files for "Rapid, Low-Cost Detection of Water Contaminants Using Regulated *In Vitro* Transcription": Handheld_Fluorescence_Illuminator_Instructions.pdf

“Rapid, Low-Cost Detection of Water Contaminants Using Regulated *In Vitro* Transcriptions”
**Alam, Jung, *et al*. (2019)**
Supplementary Data File

### Handheld Fluorescence Illuminator Instructions

3D print the following .STL files using fused filament fabrication (also known as fused deposition modeling, FDM). For the illuminators described here, we have had success using a Stratasys Fortus 250MC printer with black ABS plastic, 0.007 inch (0.178 mm) layer thickness. We have included our CAD files for anyone to develop and customize their own illuminator on top of our design.

- Front case: 786018-2_rev1-logo.STL
- Back case: 786018-3._rev1.STL
- Battery cover: 786018-4_rev1.STL
- Window frames (print 2): 786018-5 WINDOW FRAME.STL

We have also provided a printed circuit board (PCB) design for easy assembly of the electronic components. For the illuminators described here, we had our PCBs manufactured by JLBPCB.com as 2-layer boards. The files for the board can be opened using Eagle (free download at https://www.autodesk.com/products/eagle/free-download).

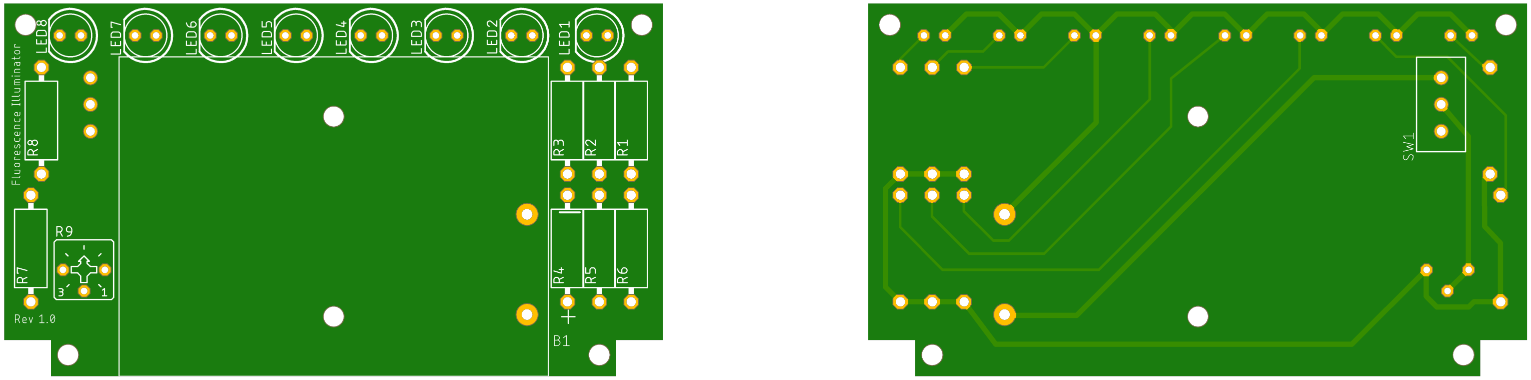


Top view of the board (left). Bottom view of the board (right). The switch (“SW1”) goes on the bottom side of the board.

Alam, Jung, *et al.* 2019.

Additional parts needed to assemble the illuminator include: 3 AAA batteries, 8 LEDs (LED1-LED8 in the illustration), 8 resistors (R1-R8), a trim potentiometer (R9) that allows you to easily adjust the brightness of the LEDs, and a switch (SW1). You will also need a set of electronics tools (T6 Torx driver, No. 1 Phillips driver, small wire clippers, etc.), a soldering iron and solder.

| Part | Supplier | Manufacturer | Supplier Part Number | Cost (April 2019) | Quantity | Details |
|---|---|---|---|---|---|---|
| 5 mm Standard Through Hole LED, 470 nm Blue | Mouser | Cree, Inc. | 941-C503BBANCZ0A0451 | $0.21/each | 8 | 15° viewing angle, 11000 mcd, $I_F$ = 20 mA, $V_F$ = 3.2 V |
| Through Hole Axial Resistor | Digi-Key | Yageo | 64.9XBK-ND | $0.10/each | 8 | 64.9 Ohms, 1/4W |
| Through Hole Trimmer Potentiometer | Digi-Key | Nidec Copal Electronics | CT6EP201-ND | $0.85 | 1 | 200 Ohm, 1/2W |
| Through Hole Slide Switch | Digi-Key | TE Connectivity ALCOSWITCH Switches | 450-1598-ND | $0.98 | 1 | |
| 3 Cell AAA Battery Holder | Digi-Key | Memory Protection Devices (MPD) | BC3AAAPC-ND | $1.94 | 1 | |
| Phillips Rounded Head Thread-Forming Screws for Plastic (#1 Phillips) | McMaster-Carr | | 99461A710 | $0.235/each (sold in pack of 50) | 6 | 18-8 Stainless Steel, Number 2 Size, 3/16" Long |
| Flat Head Thread-Forming Screws for Plastic (T6 Torx) | McMaster-Carr | | 95893A550 | $0.257/each (sold in pack of 50) | 4 | 18-8 Stainless Steel, Number 2 Size, 1/4" Long |

Assembly of the board can be tricky, especially if you're not familiar with electronics and soldering. Be prepared to make mistakes (order some extra LEDs). Although the tolerances are fairly generous, it still takes a certain level of skill to put this together. Your local research shop, core facility, maker space, etc. should have hobbyists and experts who can help you throughout this process.

**Tips:**

- Test all of your components before soldering them onto the board.
- Start with the LEDs, then add the resistors, the switch, the trim potentiometer, and finally the battery holder.
- Position the LEDs so that they first extend away from the board, and then bend 90° upwards. This is one of the challenging parts of assembling the board. We found that bending the legs around a small cylindrical rod helped to consistently position all of the LEDs. Test fit the board in the case throughout the process.
- Start with LED1, then LED8, LED2, LED7, etc. Alternating in this manner results in a more consistent build.
- The legs of the LEDs, resistors, battery holder, and switch must be trimmed as close to the board as possible. Failing to do so will make it more difficult to mount the board inside the illuminator.
- Remember that the switch mounts on the other side of the board.
- Test the board before mounting it into the illuminator.

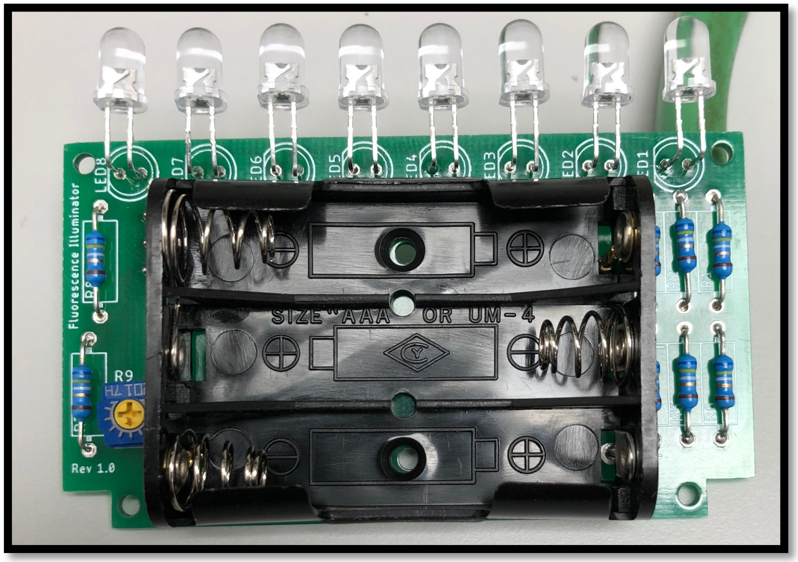


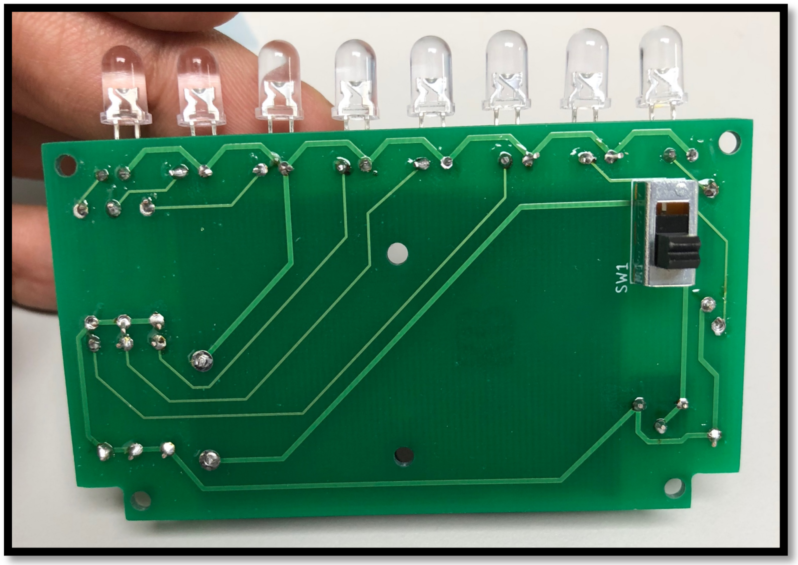


Alam, Jung, *et al.* 2019.

**Assembly of the handheld fluorescence illuminator.** (**A**) Place the front case flat on a solid surface with the backside facing towards you. (**B**) Gently place the fully-assembled printed circuit board (PCB), without batteries, into the back of the front case. Be careful to make sure that the switch on the PCB is aligned with the switch hole in the front case. Make any final adjustments (*e.g.* LED positioning) before moving onto the next step. (**C**) Place 6 round head Phillips screws into the positions shown by the arrows and secure into place with a #1 Phillips driver. (**D**) Gently place the back case onto the the front case. Do not force into position! If you are unable to place the back case on, you may need to adjust the PCB or sand down the 3D printed pieces. (**E**) Screw the back case on using 4 of the flat head screws with a T6 Torx driver. Add batteries and slide battery cover up to close.

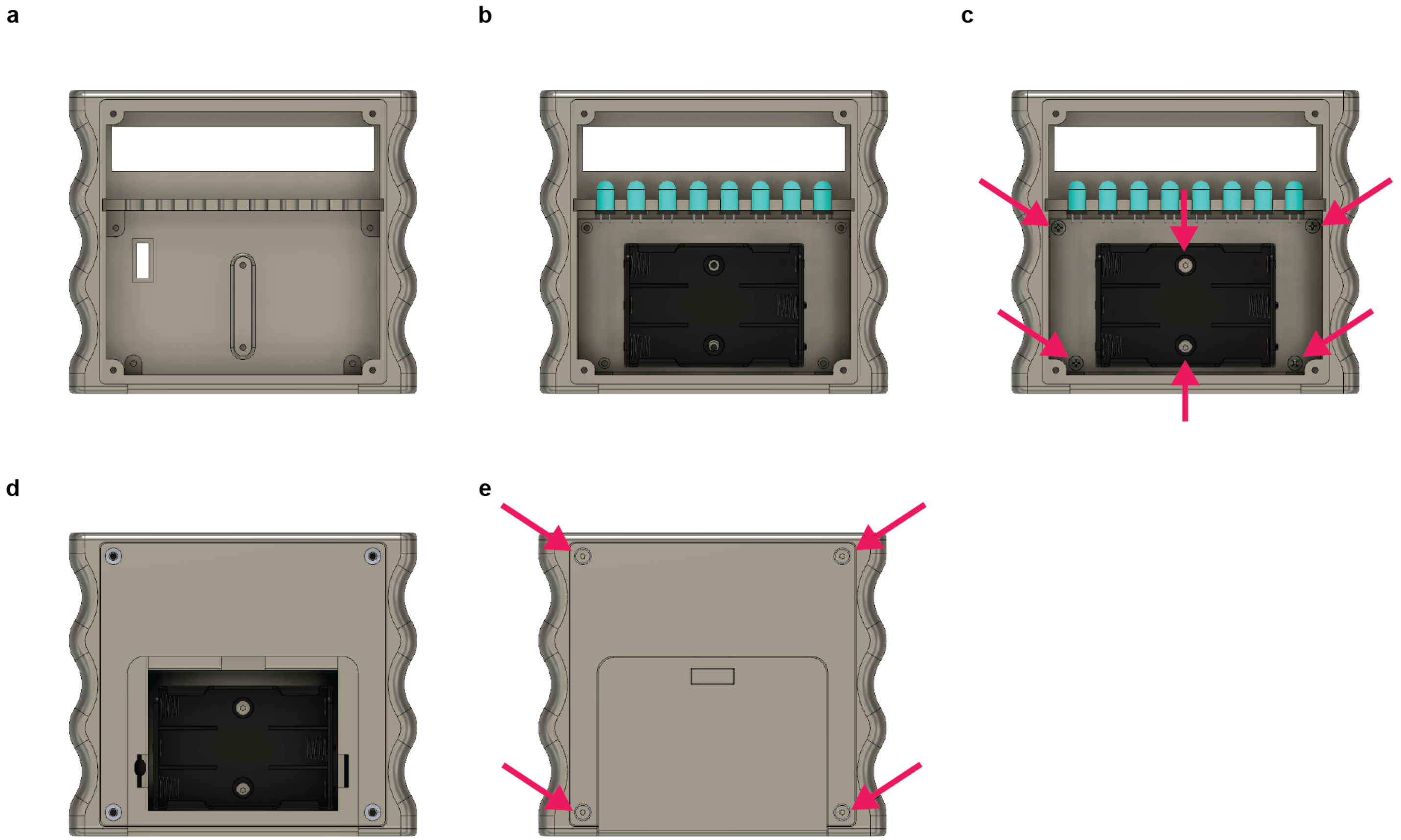


Alam, Jung, *et al.* 2019.

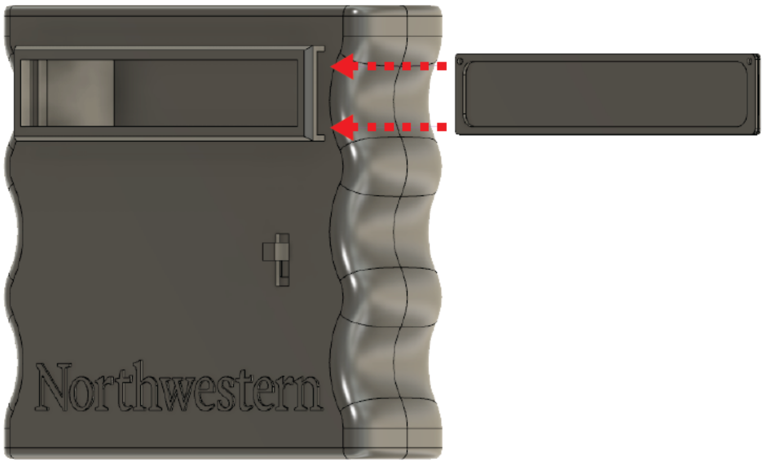


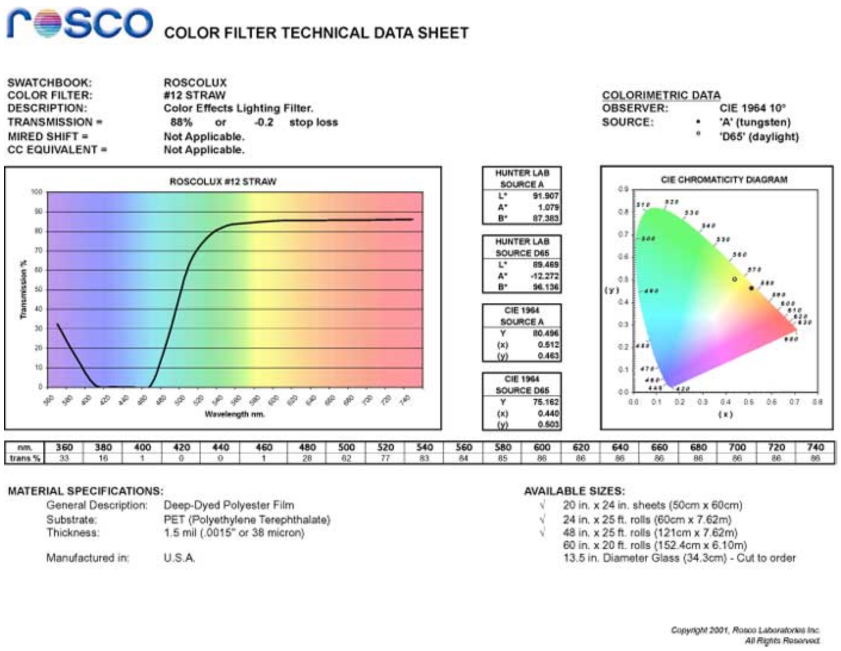

rosco COLOR FILTER TECHNICAL DATA SHEET

| | |
|---|---|
| SWATCHBOOK: | ROSCOLUX |
| COLOR FILTER: | #12 STRAW |
| DESCRIPTION: | Color Effects Lighting Filter. |
| TRANSMISSION = | 88% or -0.2 stop loss |
| MIRED SHIFT = | Not Applicable. |
| CC EQUIVALENT = | Not Applicable. |

COLORIMETRIC DATA
OBSERVER: CIE 1964 10°
SOURCE: • 'A' (tungsten)
° 'D65' (daylight)



| HUNTER LAB SOURCE A | |
|---|---|
| L* | 91.907 |
| A* | 1.079 |
| B* | 87.383 |

| HUNTER LAB SOURCE D65 | |
|---|---|
| L* | 89.469 |
| A* | -12.272 |
| B* | 96.136 |

| CIE 1964 SOURCE A | |
|---|---|
| Y | 80.496 |
| (x) | 0.512 |
| (y) | 0.463 |

| CIE 1964 SOURCE D65 | |
|---|---|
| Y | 75.162 |
| (x) | 0.440 |
| (y) | 0.503 |

| nm. | 360 | 380 | 400 | 420 | 440 | 460 | 480 | 500 | 520 | 540 | 560 | 580 | 600 | 620 | 640 | 660 | 680 | 700 | 720 | 740 |
|---|---|---|---|---|---|---|---|---|---|---|---|---|---|---|---|---|---|---|---|---|
| trans % | 33 | 16 | 1 | 0 | 0 | 1 | 28 | 62 | 77 | 83 | 84 | 85 | 86 | 86 | 86 | 86 | 86 | 86 | 86 | 86 |

MATERIAL SPECIFICATIONS:
General Description: Deep-Dyed Polyester Film
Substrate: PET (Polyethylene Terephthalate)
Thickness: 1.5 mil (.0015" or 38 micron)

Manufactured in: U.S.A.

AVAILABLE SIZES:
√ 20 in. x 24 in. sheets (50cm x 60cm)
√ 24 in. x 25 ft. rolls (60cm x 7.62m)
√ 48 in. x 25 ft. rolls (121cm x 7.62m)
60 in. x 20 ft. rolls (152.4cm x 6.10m)
13.5 in. Diameter Glass (34.3cm) - Cut to order

Copyright 2001, Rosco Laboratories Inc.
All Rights Reserved.

Alam, Jung, *et al.* 2019.

The window assembly consists of two identical frames, so make sure you print two of them. They are designed to sandwich a piece of stage light film between them and slide as a single unit into the window on the front case of the illuminator. Test fit the frames and adjust as needed before gluing the filter and frames together. We recommend using a small amount of a strong and clear adhesive (*e.g.* cyanoacrylate, a.k.a. “superglue”). Once assembled, the framed filter should easily slide into the window.

For visualizing green fluorescence, we recommend using Roscolux #12 Straw as a lighting filter. A poster size roll can be purchased online for ~$10 (before shipping and handling). It efficiently transmits light at wavelengths longer than 500 nm, while blocking a sufficient amount of blue light across 380–480 nm. This makes it a practical solution for blocking the blue light generated by the LEDs, but not the green fluorescence generated in ROSALIND reactions. An additional layer of film can improve the blue light blocking but will also result in lower transmittance of green light.

A benefit of having a removable window is that you can easily try several different filters until you find something that works best for your application.

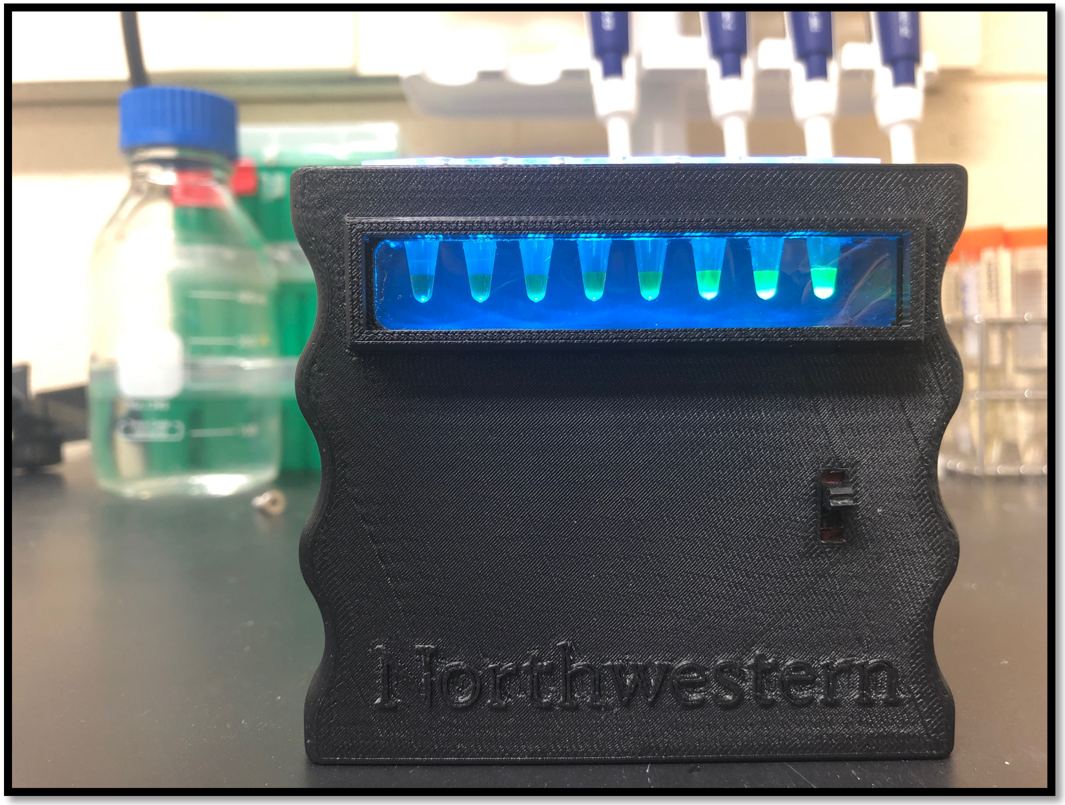


Contact the Lucks Lab (http://luckslab.org) with any questions. Post and share your builds with us by tagging us on Twitter @LucksLab!

**Acknowledgements:** We acknowledge the work of Northwestern University's Research Shop for the CAD designs and development of the PCB, and Northwestern's Rapid Prototyping facility for manufacturing them.
