## Supplementary figures and images for "Rapid, Low-Cost Detection of Water Contaminants Using Regulated *In Vitro* Transcription"

### Fig.S7_IMG1.JPG

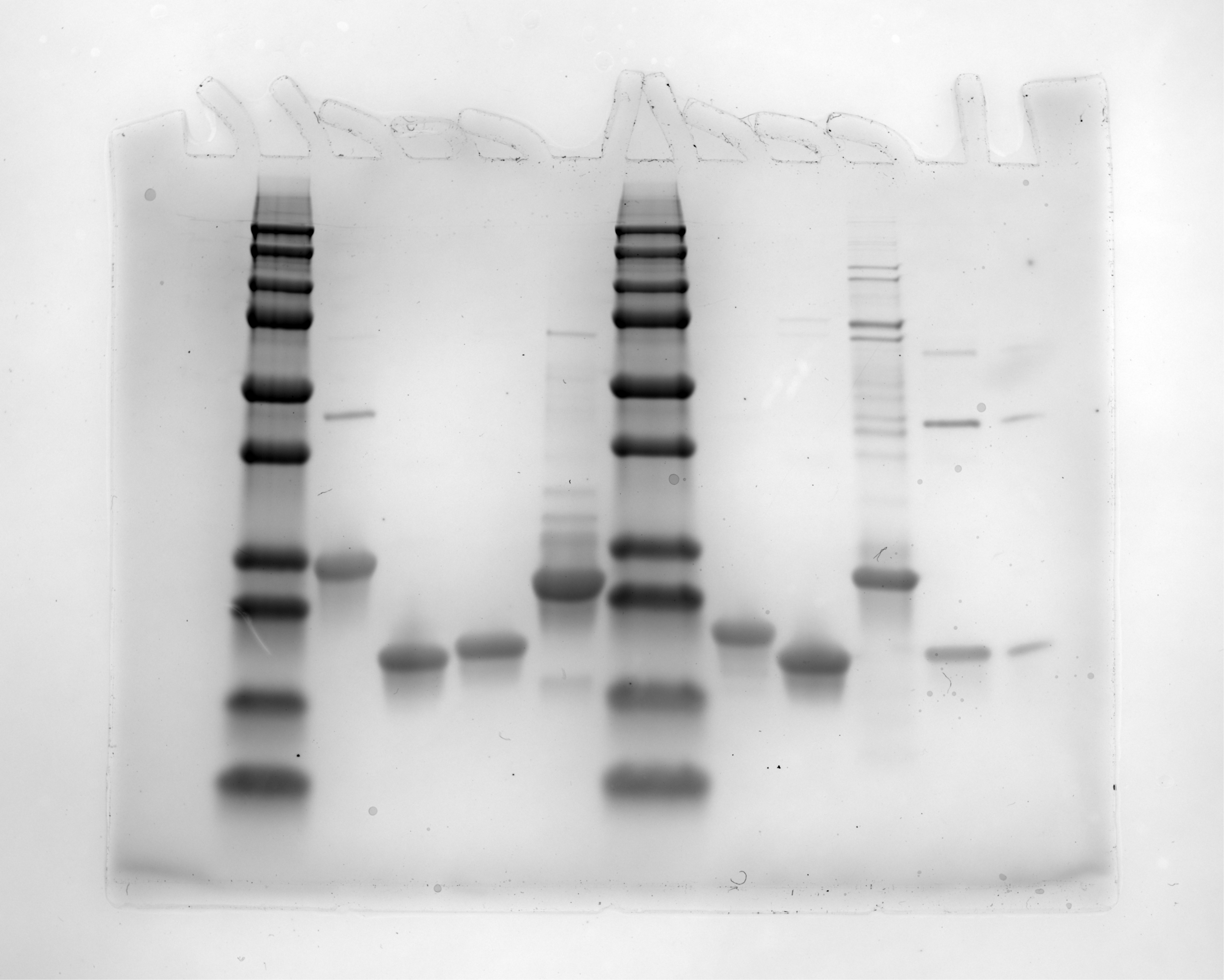

### Fig.S7_IMG2.JPG

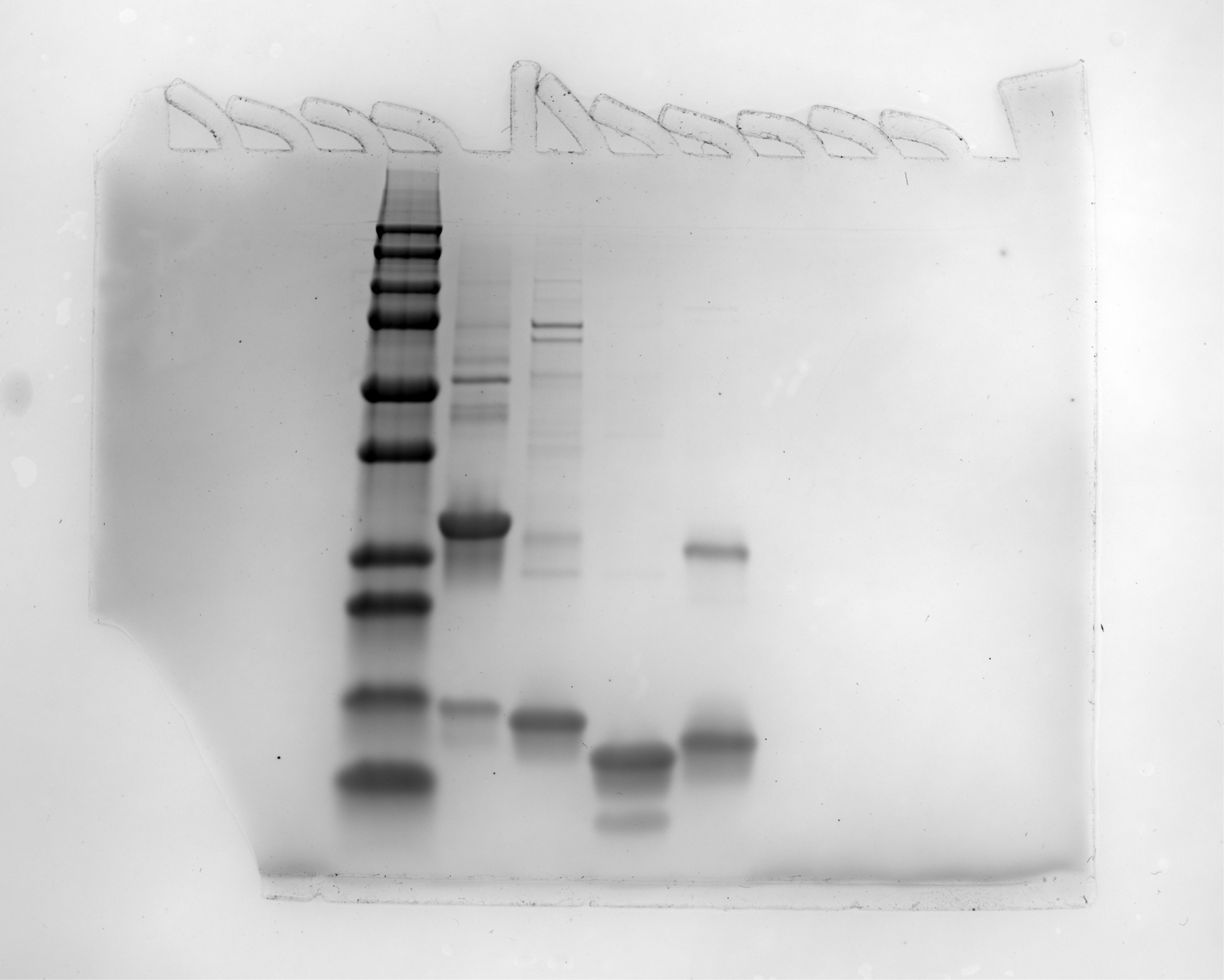

### IMG_1360.JPG

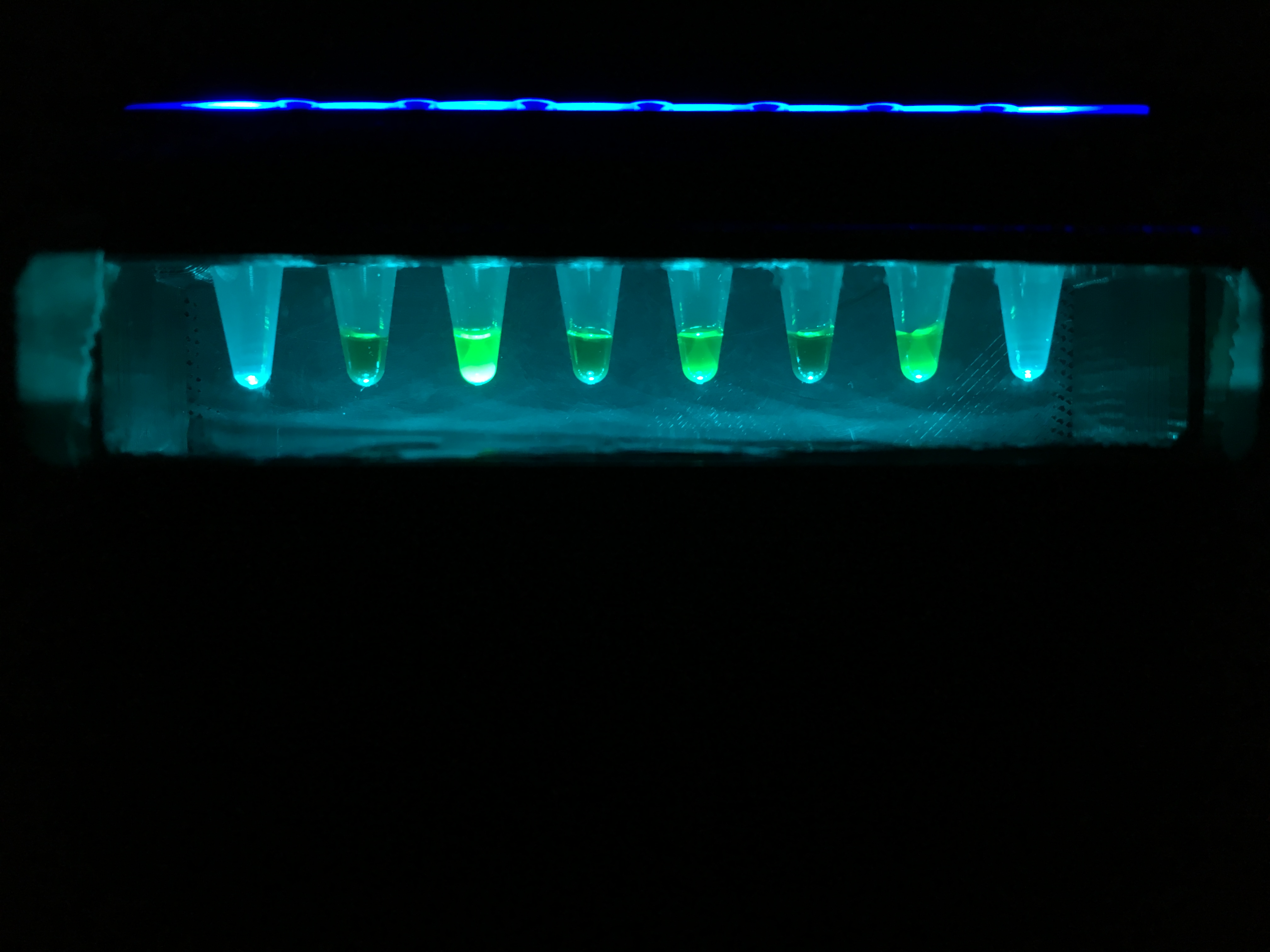

### IMG_1364.JPG

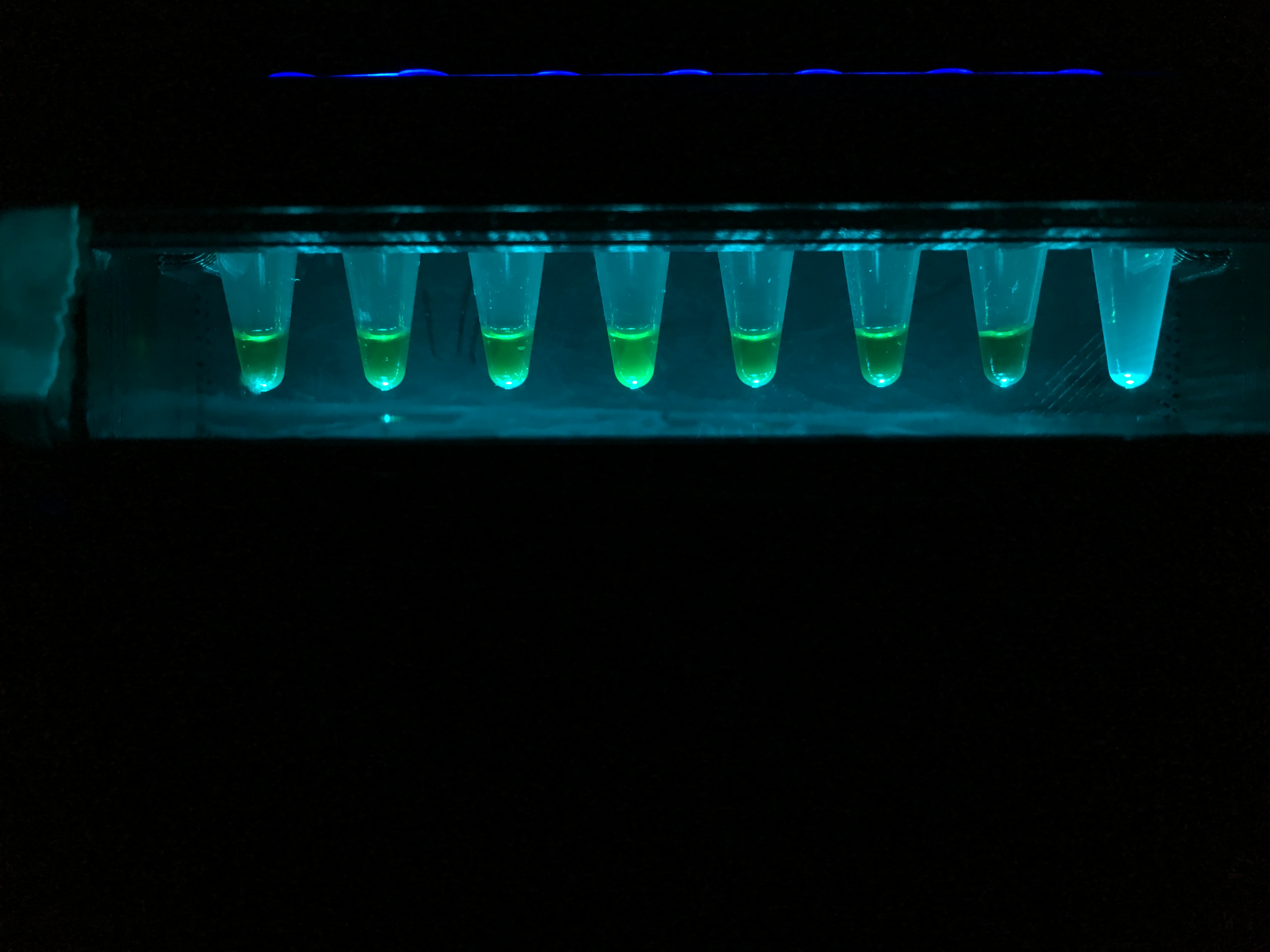

### IMG_1408.JPG

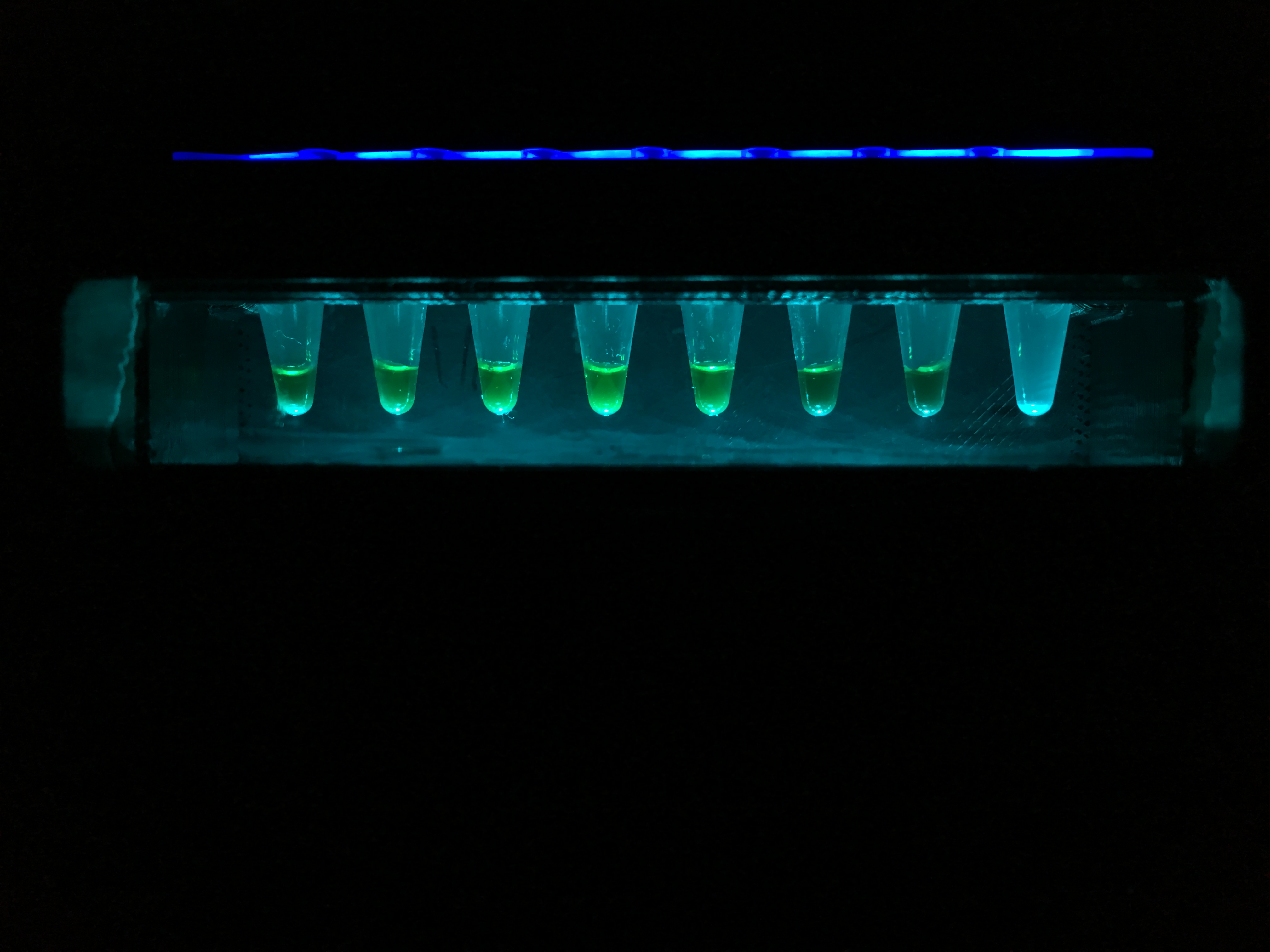

### IMG_1425.JPG

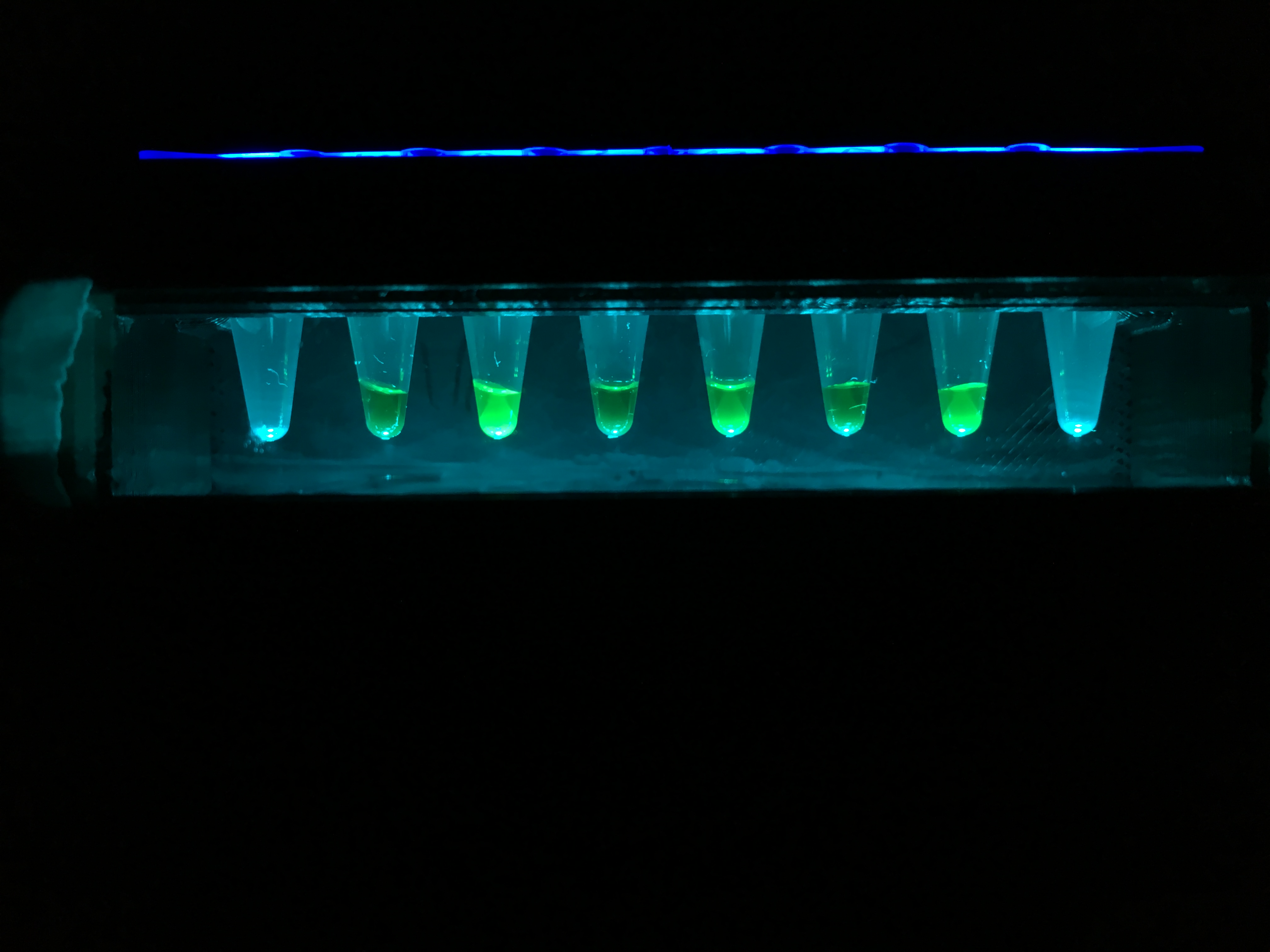

### IMG_5546.JPG

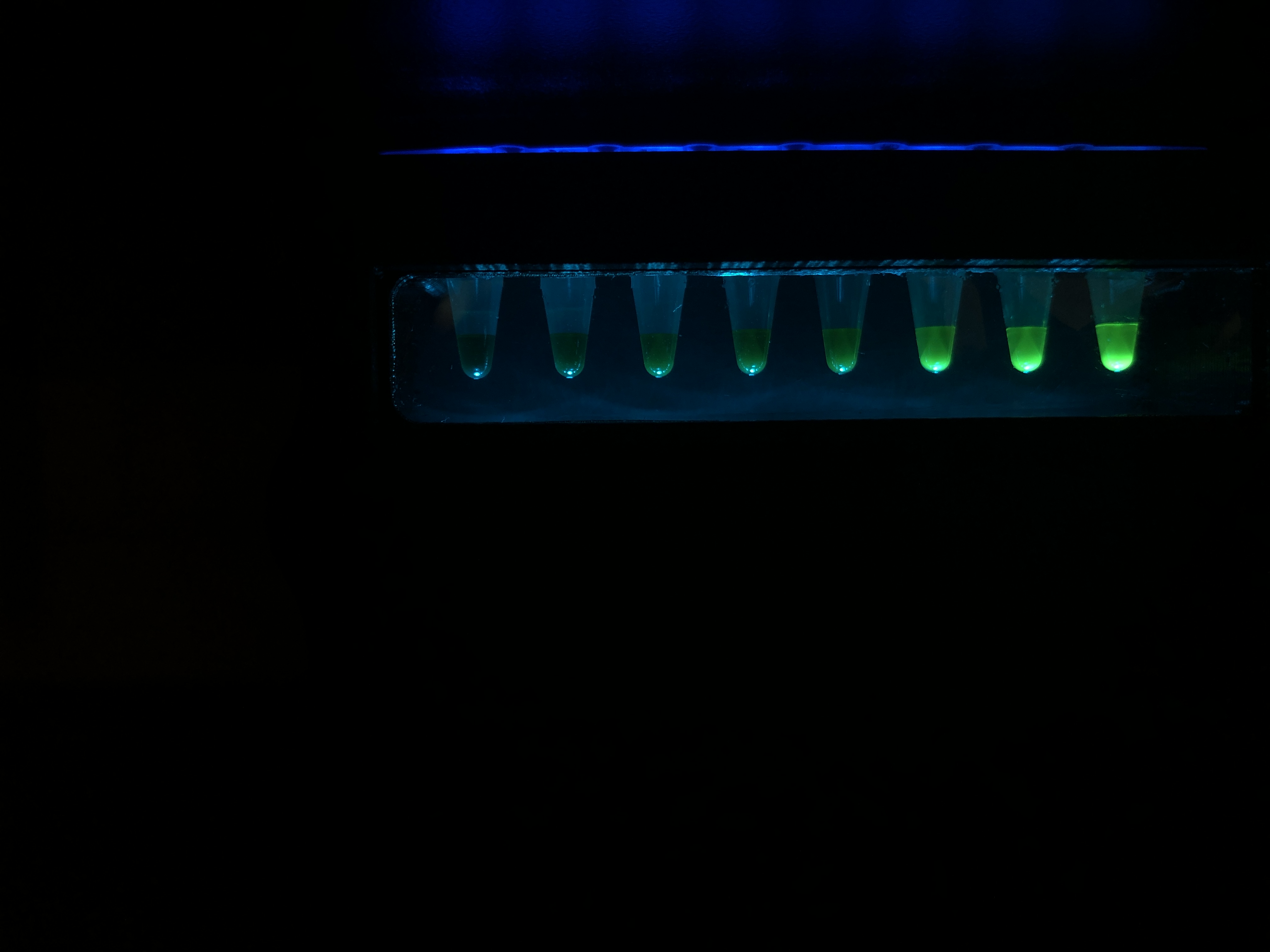
